# Farming and climate legacies shape the seed microbiota and offspring drought responses in wheat

**DOI:** 10.64898/2026.08.08.743720

**Authors:** Barkha Sharma, Josephine Burgmans, Niels N. Oehlmann, Johannes G. Rebelein, Martin Schädler, Hamed Azarbad

## Abstract

Seeds link parental environments to offspring establishment, but whether seed-associated bacteria retain signatures of farming and climate legacies across plant generations remains unclear. Here, we characterized epiphytic and endophytic bacterial communities of winter wheat seeds collected from the Global Change Experimental Facility (GCEF) in Germany across three harvest years representing contrasting climates. We then tested how farming (organic versus conventional) and climate (ambient versus future) legacies experienced by maternal plants were associated with offspring rhizosphere bacterial communities and plant performance under drought in the greenhouse. Harvest year was the dominant driver of grain dry weight and seed-associated bacterial communities. Climate legacy additionally affected seed epiphytic communities, whereas farming legacy was expressed in the endophytic diversity. Germination was higher overall for seeds from the conventional than the organic farming legacy and from the ambient than the future climate legacy. A small subset of unique seed-associated ASVs was detected in offspring rhizospheres. Although these ASVs occurred at low relative abundance in seeds (<1%), they accounted for up to approximately 40% of rhizosphere relative abundance under drought. Together, these findings show that seeds retain bacterial signatures of parental farming and climate legacies and that a subset of seed-associated ASVs remains detectable and can become abundant in offspring rhizospheres under drought.

## 1. Introduction

Wheat (*Triticum aestivum* L.) is cultivated on more than 220 million hectares worldwide and provides nearly one-fifth of all calories and protein consumed by humans, making it central to global food security [1–3]. This role is under mounting threat from the climate. In Europe, the 2018 growing season brought one of the most severe compound drought-heat events on record, causing widespread yield losses [4,5], highlighting the vulnerability of crop production to increasingly frequent climatic extremes [6]. At the same time, European agriculture is changing. Organic farming has expanded threefold over the past two decades, driven by environmental policy and consumer demand [7,8]. This shift changes not only the chemistry and biology of managed soils but also the microbiological environment in which crops develop [9]. A central question is therefore how intensifying climate variability and diverging farming systems interact to shape wheat crop biology, and whether their effects persist beyond the seasons of exposure.

Plant stress responses are shaped not only by host genetics but also by associated microbial communities [10,11]. As an example, root-associated microbiomes can improve water-use efficiency, produce osmolytes and phytohormones that buffer osmotic stress, and shift towards stress-tolerant taxa under drought [12–14]. Yet these microbial communities are often treated as primarily soil-derived i.e., assembled anew each generation from the local edaphic pool [15]. This view is being revised by evidence that seeds harbour bacterial communities that differ from bulk soil in composition and ecological origin, and (part of) seed-associated microbes can contribute to the microbiota of emerging seedlings during germination and early growth [16–18]. This contribution of seed-associated microbes provides a potential route through which parental environmental history may remain detectable in the next generation [19], allowing seeds to act as potential carriers of microbial function [20,21]. This process forms the basis of microbial memory, whereby past environmental exposure may shape future microbial response through legacy or history effects. Such legacy effects may influence plant and microbial responses to subsequent stress, but their ecological relevance across plant generations remains poorly understood [22].

Seed microbiota are shaped mainly by two distinct routes. (i) Vertical transmission (transfer from maternal plant to its offspring) via maternal tissues such as flowers, ovules, or embryos [23,24], which may support the persistence of a subset of seed-associated taxa across plant generations [25,26]. (ii) Horizontal acquisition (recruitment from the surrounding environmental reservoirs) during seed development, dispersal or germination [27,28], which may increase microbiome responsiveness to local conditions [29,30]. These routes contribute differently to bacterial communities located inside and on the surface of seeds. Endophytes (inside seed tissues) are often more persistent across generations and may be more strongly influenced by host filtering and parental transmission [31–33]. During germination, some seed endophytes can colonize emerging roots, resulting in priority effects, meaning that early colonists influence the establishment of later-arriving microorganisms [18,34]. Epiphytic communities (external seed surface) are more directly exposed to environmental filtering [35,36]. Despite increasing evidence that seed-associated microbes contribute to early plant microbiota [37], the environmental legacies that structure these communities and determine their relevance to the next generation remain poorly understood [19].

Drought can restructure wheat-associated rhizosphere and phyllosphere microbiomes within a single growing season, partly through changes in root exudate profiles that favour stress-tolerant taxa [38]. Because seed microbiota are assembled during anthesis and grain filling, stages highly sensitive to water limitation, climate conditions during seed development can act as a temporal filter on seed microbial composition, with potential consequences for the microbiota and stress responses of the next generation [22,39]. Agricultural management imposes an additional filtering layer [40,41]. It has been shown that long-term organic farming supports greater soil microbial diversity, network complexity, and functional potential than conventionally managed soils [42–45]. These soil microbial differences can further propagate into root and shoot microbiomes through vascular transmission during seed development [46]. However, it remains unclear how contrasting climate and farming histories leave distinct signatures in seed epiphytes and endophytes and whether these signatures are associated with offspring performance under drought.

Here, we used winter wheat seeds collected from the Global Change Experimental Facility (GCEF), Germany, a long-term field experiment where organic and conventional farming systems were maintained under ambient and future climate scenarios [6]. Seeds were harvested in three contrasting harvest years, ranging from extreme drought in 2018 to intermediate conditions in 2021 and comparatively wet conditions in 2024. We characterized epiphytic and endophytic seed bacterial communities and tested how climate and farming legacy were associated with offspring performance and rhizosphere bacteria under well-watered and drought conditions in the greenhouse. We hypothesized that (i) harvest year would be the dominant driver of seed bacterial diversity and composition, with the 2018 extreme drought imposing the strongest compositional filtering across seed compartments; (ii) climate- and farming legacies would leave compartment-specific signatures in seed bacterial communities, and a subset of seed-associated ASVs would also be detected in offspring rhizospheres; and (iii) these legacy-associated microbial patterns would be associated with variation in offspring germination, growth and physiological responses to drought.

## 2. Materials and Methods

### 2.1. Field experiment in GCEF, Halle

The field experiment was conducted at GCEF, a large multi-year field experiment in Saxony-Anhalt, Germany (51° 23’ 30N, 11° 52’ 49E, 116 m a.s.l.), which is designed to simulate the changing global climate conditions to examine the effect of different ecosystem processes across various land-use types and intensities (Fig. 1A). The field consisted of 50 plots (16 m × 24 m each) arranged into 10 main plots, with 5 sub-plots per main plot [6]. The seeds were collected from plot under two types of land-use regime: (a) conventional farming: mineral fertilizers and pesticides applied as per conventional agricultural practices, (b) organic farming: mechanical weed control, organic fertilization, untreated seeds, and pesticide are not used (Fig. 1A). In conventional farming, the crop rotation consists of winter rape, winter wheat, and winter barley, with the use of synthetic fertilizers and pesticides (see Schädler et al. (2019); Sharma et al. (2026) [6,47] for detailed information). To simulate future climate conditions, half of the experimental blocks (5 blocks) were placed under a steel structure equipped with a mobile roof, side panels, and an irrigation system (Fig. 1A). By closing the roof and side panels, night temperatures were elevated, and summer precipitation was reduced by approximately 20%, while spring and autumn precipitation were increased by 10% via irrigation. The other 5 blocks, used as controls, were housed under a similar structure without interventions to maintain ambient climate conditions. Air temperature and precipitation data were obtained from the on-site meteorological monitoring system at GCEF. Air temperature (°C) was recorded daily at a height of 30 cm above the soil surface within each subplot. Precipitation was measured once per main plot (block) and recorded as the daily cumulative sum (mm, equivalent to Lm^-2^).

**Figure 1.**
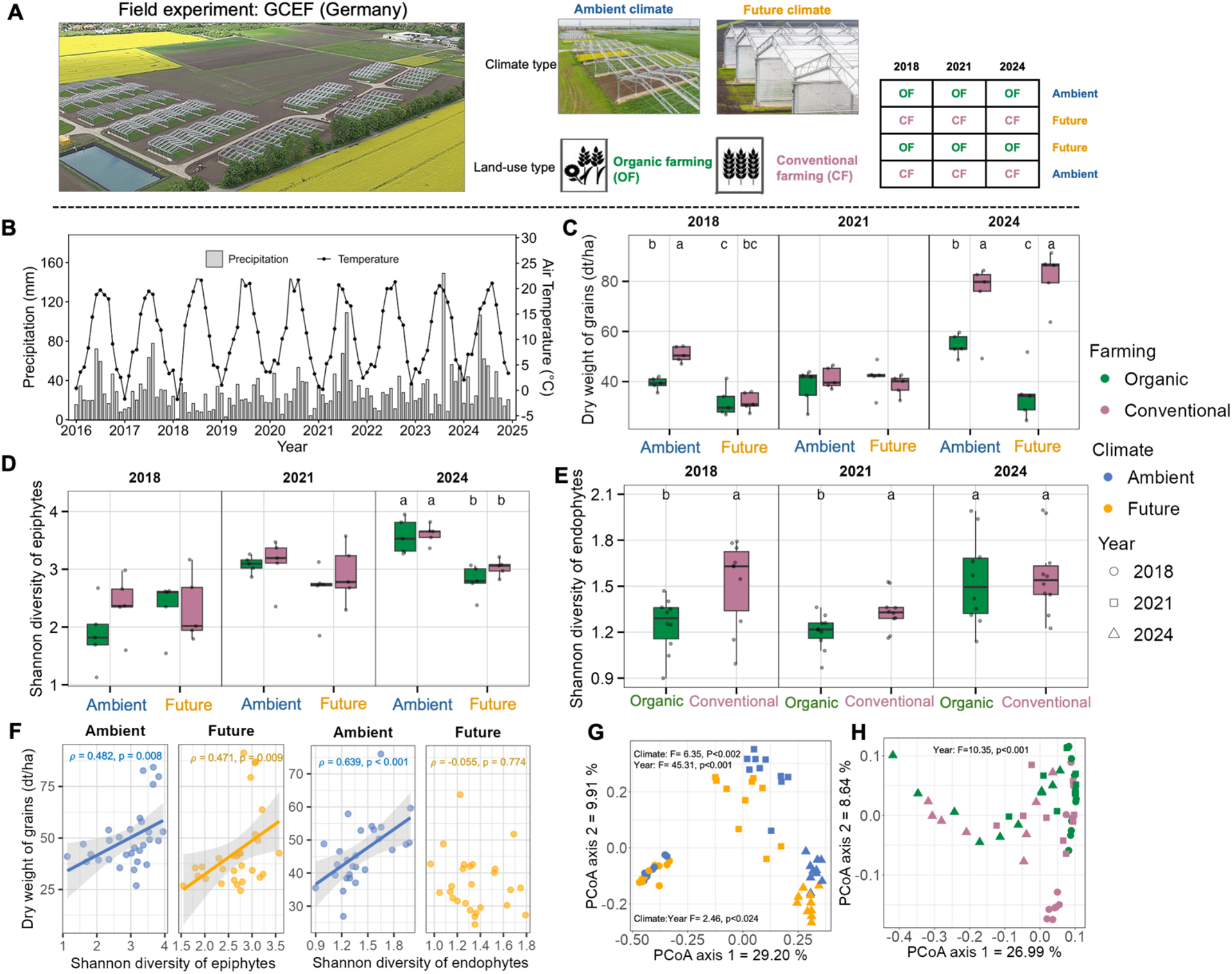
Climatic context, seed traits and bacterial communities across farming and climate legacies. (A) Aerial view and the experimental design of Global Change Experimental Facility (GCEF) experiment at Bad Lauchstädt, Germany. Winter wheat (*Triticum aestivum* L. cv. RGT Reform) was grown under two farming systems (organic farming (OF) and conventional farming (CF)) and two climate treatments (ambient and future climate). Seeds harvested in 2018, 2021 and 2024 were used in the present study. (B) Monthly precipitation and mean air temperature at the GCEF from 2016 to 2024. (C) Dry weight of harvested winter wheat grain under the four farming × climate legacy combinations in 2018, 2021 and 2024. Different letters in the panel denote statistically significant differences based on post hoc Tukey’s test (*P*<0.05). (D-E) Shannon diversity of seed epiphytic (D) and endophytic seed bacterial communities. (F) Associations between grain dry weight and Shannon diversity of epiphytic and endophytic seed bacterial communities under ambient and future climate legacies. Lines and shaded areas show fitted linear trends and 95% confidence intervals for visualization. Spearman’s correlation coefficients (ρ) and Benjamini–Hochberg-adjusted P values are shown in each panel. (G) Principal coordinate analysis (PCoA) based on Bray-Curtis dissimilarities of seed epiphytic (G) and endophytic (H) seed bacterial community composition. Samples are coloured by climate legacy in G and farming legacy in H, and symbols denote harvest year. PERMANOVA results for significant effects are shown within each panel.

We used winter wheat seeds (*Triticum aestivum* L., cv. RGT Reform) collected from the GCEF during three growing seasons (2018, 2021, and 2024). When evaluating effects on harvested seeds and offspring plants, parental treatments are hereafter referred to as farming legacy and climate legacy, respectively. Consistent with the respective farming systems, organic plots were sown with untreated seeds (T0), whereas conventional plots were sown with fungicide-coated seed. Each legacy combination was represented by five replicate plots, resulting in 20 plots per harvest year and 60 plot-year seed batches across the three harvest years. Grain dry weight was recorded for each treatment after threshing. All harvested seed samples were stored at room temperature (15-18°C) until further investigation. Consequently, seed batches harvested in 2018, 2021 and 2024 differed in storage duration. However, because all farming and climate legacy treatments within a given harvest year experienced the same storage duration, within-year analyses allowed these legacy effects to be assessed without direct confounding by differences in storage duration among harvest years. Details on extraction of the seed endophytes and epiphytes are presented in detail in the Supplementary material and methods.

### 2.2. Greenhouse experiment

To study the impact of farming and climate legacies of seed microbiota on next-generation plants, a pot experiment was conducted in the greenhouse at Marburg University (50.8083° N, 8.7694° E). Seeds harvested from the field experiment were sown in potting soil with four seeds planted per pot at a depth of 2 cm. Plants were grown under Neusius LED grow lights (Neusius Pflanzenlicht, Item-Nr. LED/D-E4-180VR) with a 12-hour light/dark photoperiod in a greenhouse (Fig. 3A). The pots were watered with 100 ml of tap water every two days for two weeks to ensure consistent initial growth conditions. Germination rate and time were measured over a period of 6 days. Germination rate was calculated as the percentage of seeds that successfully sprouted each day relative to the total number of seeds per pot (4 seeds). After 14 days of seedlings established, watering conditions were adjusted, where water stress (drought) was then applied to half of the pots, with drought-treated pots receiving 40 ml of water compared to 100 ml for well-watered control plants. The watering schedule was adjusted based on temperature and upper soil moisture to maintain drought conditions for six more weeks. Pots were arranged in a fully randomized design, and their positions were re-randomized weekly. The experimental setup resulted in the total combination of the following treatments: 60 [2 farming (organic vs. conventional) × 2 climate (ambient vs. future) × 3 years (2018, 2021, and 2024) × 5 field sub-plot replicates] × 2 water stress treatments (drought vs. well water control). Each treatment was replicated twice in a randomized design (120 treatments × 2 replicates = 240 pots).

Plant height and chlorophyll content (using Chlorophyll meter SPAD 502 plus) were measured at two time points: before drought exposure (2 weeks of the experiment) and at harvest (after 6 weeks of drought exposure). Aboveground biomass was harvested to determine fresh and dry weight. Dry biomass was obtained after oven-drying samples at 60 °C for 4 days, until a constant weight was achieved. Dry weight content (DWC) was calculated as the ratio of dry to fresh biomass and expressed as a percentage. Rhizosphere soil was collected at harvest by carefully uprooting plants and removing loosely attached soil [9]. Details on the mesurment of the rhizosphere CO_2_ production and rhizosphere DNA extraction, amplicon library preparation, sequencing, and bioinformatic analyses are presented in detail in the Supplementary material and methods.

### 2.3. Statistical analysis

All statistical analyses were conducted in R (Version 4.3.2; R Development Core Team, 2024) using R Studio (Version 2023.12.1+402). Climate variables at GCEF (air temperature and precipitation) were analyzed using linear-mixed models (LMMs) with harvest year, month, and their interactions were included as fixed effects, while field subplot was included as a random intercept. Unless otherwise stated, response variables measured across the three harvest years (e.g., grain dry weight and α-diversities) were first analysed using ANOVA that included harvest year and the relevant experimental factors and their interactions. Separate models were then fitted within each harvest year to characterize year-specific effects of farming and climate legacies and, for the greenhouse experiment, watering treatment. Within-year analyses compared seed batches stored for the same duration and therefore allowed farming legacy and climate legacy effects to be evaluated without direct confounding by differences in storage duration among years. Germination data were analysed using a binomial generalized linear mixed model (GLMM) with farming legacy, climate legacy, and harvest year as fixed factors, followed by Tukey-adjusted pairwise comparison of estimated marginal means. Plant height, fresh biomass, DWC, chlorophyll content (SPAD index), and rhizosphere respiration (CO_2_) were analysed using factorial ANOVA models to evaluate variations in plant performance across experimental factors, including farming legacy, climate legacy, harvest year, and water stress, followed by Tukey’s HSD tests. Shannon diversity was used to characterize bacterial α-diversity. Community structure was assessed using Bray–Curtis dissimilarities, visualized by principal-coordinate analysis and tested by PERMANOVA. Spearman correlations were used to assess relationships among seed traits, microbial diversity and plant traits, with Benjamini– Hochberg correction for multiple testing.

ASVs unique to or shared among rhizosphere communities under well-watered and drought conditions were summarized separately for each harvest. To assess bacterial ASV overlap between seeds and offspring rhizospheres, ASV occurrence and relative abundance were compared separately between epiphytic or endophytic seed bacterial communities and drought-treated rhizospheres within each harvest year and farming legacy. For these analyses, ASVs were classified into four categories: (i) Shared, representing ASVs detected in both seed and offspring rhizosphere across harvest years; (ii) Unique, representing harvest year-specific ASVs detected in both seeds and the corresponding offspring rhizosphere; (iii) Lost, representing seed-associated ASVs that were not detected in the offspring rhizosphere; and (iv) Acquired, representing ASVs detected in the offspring rhizosphere but not in the corresponding seed samples. Full statistical procedures, model specifications, taxon-level analyses and R packages are provided in Supplementary Methods.

## 3. Results

### 3.1. GCEF field experiment

#### 3.1.1. Climate context and grain dry weight

To characterize the parental environmental context of the sampled seeds, we used winter wheat collected from the GCEF in 2018, 2021, and 2024 (Fig. 1A). These harvest years represented contrasting climatic histories at the study site (Fig. 1B; Table S1). During the growing season (April–September), air temperature and precipitation differed significantly among harvest years (temperature: F = 633.3, P < 0.001; precipitation: F = 11.143, P < 0.001). Both variables also showed significant month by harvest year interactions (temperature: F = 261.78, P < 0.001; precipitation: F = 8.270, P < 0.001; Table S1). Relative to the 2016–2017 growing-season mean, 2018 was +2.2 °C warmer and received approximately 80% less precipitation. By contrast, 2021 showed intermediate conditions, with lower precipitation (∼33%) and slightly lower temperature (−0.4 °C), whereas 2024 was comparatively wetter than both 2018 and 2021 (Table S1, Fig. 1B).

Grain dry weight differed among harvest years (F = 69.08, P < 0.001), with the highest values observed in 2024 (Fig. 1C, Table S2). Within-year analyses showed that farming legacy affected grain dry weight in 2018 and 2024, but not in 2021 (2018: F = 33.60, P < 0.001; 2024: F = 56.90, P < 0.001). In 2018, grain dry weight was highest in the conventional-ambient treatment and lowest in the organic-future treatment (P < 0.001). In 2024, both conventional treatments produced heavier seeds than the corresponding organic farming legacies. Within organic farming, grain dry weight remained higher under ambient than future climate, whereas the two conventional climate treatments did not differ (Farming × Climate, 2024: F = 9.152, P = 0.016; Fig. 1C, Table S3).

#### 3.1.2. Diversity of seed epiphytes and endophyte bacteria

Seed bacterial Shannon diversity varied significantly among harvest years in both epiphytes and endophytes (epiphytes: F = 32.22, P < 0.001; endophytes: F = 8.855, P = 0.001; Table S4). Epiphytic diversity also differed significantly between climate legacies (F = 6.120, P = 0.017), and their responses depended on the harvest year (F = 5.798, P = 0.006). Within-year analyses showed that the climate legacy effect was restricted to 2024, where epiphytic Shannon diversity was significantly higher under ambient than future climate across both farming systems (F = 43.12, P < 0.001; Table S5; Fig. 1D). Endophytic diversity, by contrast, was shaped by farming (F = 6.961, P = 0.011), with higher Shannon diversity in conventionally than organically managed seeds in 2018 (F = 6.001, P = 0.026) and 2021 (F = 6.375, P = 0.023), but not in 2024 (Table S5; Fig. 1E). Under ambient climate, grain dry weight was positively correlated with both epiphytic (ρ = 0.482, p = 0.008) and endophytic Shannon diversity (ρ = 0.639, p < 0.001), whereas under future climate this positive association persisted only for epiphytes (ρ = 0.471, p = 0.009). Across harvest years, grain dry weight and overall seed bacterial diversity were negatively correlated with temperature and positively correlated with precipitation (Fig. S2).

#### 3.1.3. Seed bacterial community structure and composition

PERMANOVA showed that harvest year significantly influenced the structure of both epiphytic (F = 45.30, P = 0.001) and endophytic (F = 10.35, P = 0.001) seed-associated bacterial communities (Fig. 1G, H; Table S6). For epiphytes, community structure also differed significantly between climate legacies (F = 6.350, P = 0.002), and this effect depended on harvest year, as indicated by a significant climate × year interaction (F = 2.464, P = 0.024).

Within-year analyses identified a significant farming × climate interactions in 2018 (F = 2.012, P = 0.021) and significant climate effects in 2021 (F = 3.232, P = 0.006) and 2024 (F = 4.973, P = 0.001; Fig. S3A, Table S7). By contrast, endophytic community structure showed no overall effects of farming or climate across harvest years. However, within-year analysis identified a significant effect of farming legacy in 2018 (F = 5.316, P = 0.004; Table S7, Fig. S3B). The relative-abundance profiles of dominant bacterial groups also differed significantly among harvest years (Tables S8-9; Fig. S4). These differences were primarily observed in epiphytic communities, which showed more frequent responses to farming and climate legacies than endophytic communities (Fig. S4, Table S8, S9).

ASV occurrence patterns differed among epiphytic and endophytic seed communities across harvest year (Fig. 2). The largest compartment-year specific fraction comprised 1,060 ASVs detected only in epiphytes from 2018, followed by 226 ASVs detected only in endophytes from 2024. Across all three harvest years, 154 ASVs were consistently detected in epiphytic communities, whereas 168 ASVs were consistently detected in endophytic communities, which represent the core compartment specific ASVs. These cross-year compartment-specific ASV sets showed distinct taxonomic profiles. Endophyte-specific ASVs were dominated by *unclassified_Alphaproteobacteria*, whereas epiphyte-specific ASVs were taxonomically more diverse and included substantial contributions from *Pantoea*, *Massilia*, *Paenibacillus,* and *Pseudomonas* (Fig. 2).

**Figure 2.**
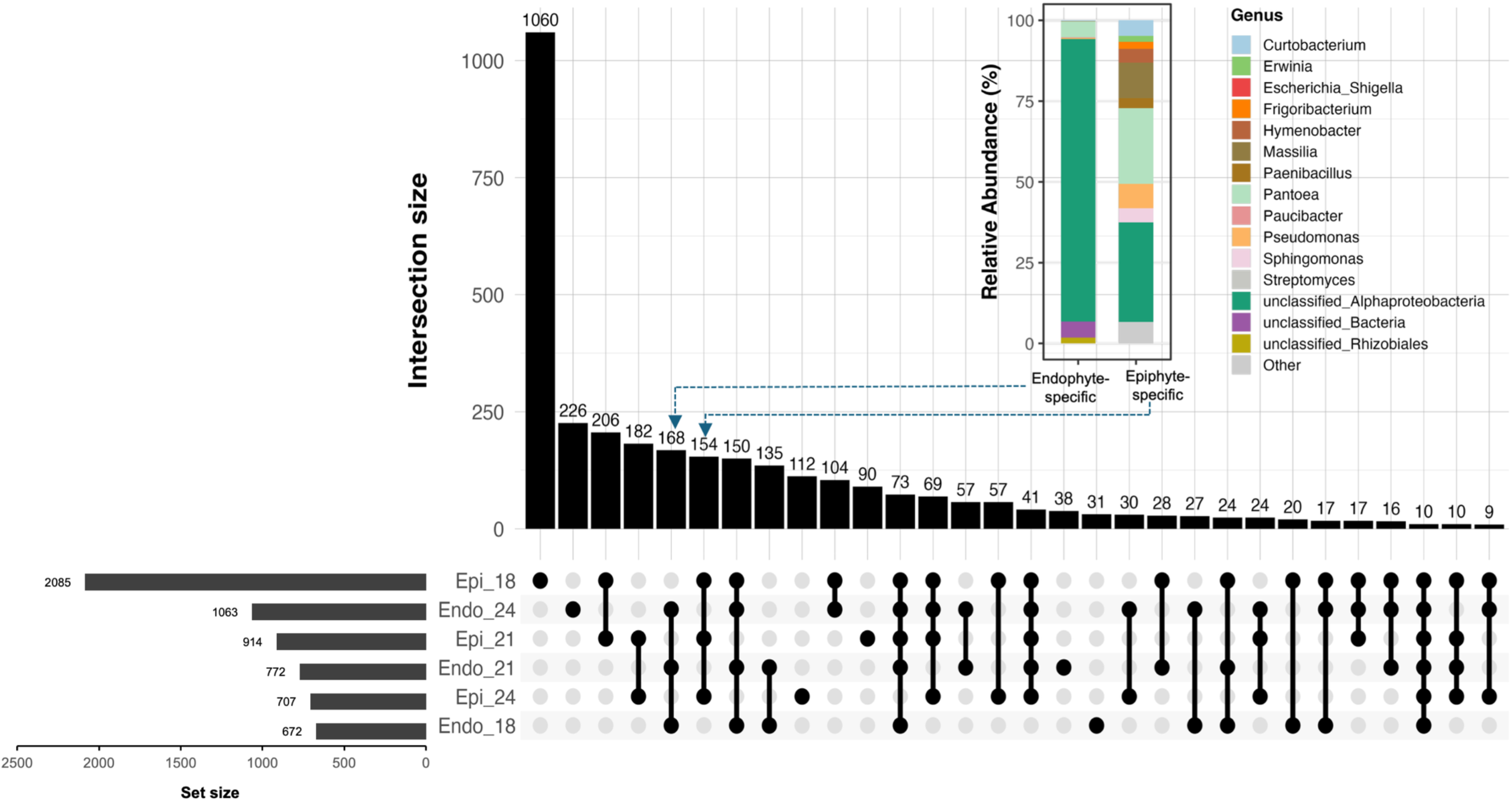
Compartment- and harvest-year-specific occurrence patterns of seed bacterial ASVs. Upset plot showing ASVs intersections among seed epiphyte and endophyte bacterial communities 2018, 2021, and 2024 harvest years. Horizontal set size bars show the total ASVs detected in each compartment–harvest-year group, whereas vertical bars show the number of ASVs in each intersection. Filled circles identify the groups included in each intersection, and connecting lines indicate intersections involving more than one group. Columns containing a single filled circle represent ASVs detected only in that compartment–harvest-year group. The inset shows the taxonomic composition, based on relative abundance, of the 168 ASVs detected in endophytic communities across all three harvest years, and the 154 ASVs detected in epiphytic communities across all three harvest years.

### 3.2. Greenhouse experiment

#### 3.2.1. Offspring establishment, growth, and physiological responses under drought

Germination was shaped by farming legacy (χ² = 20.569, P < 0.001), climate legacy (χ² = 12.912, P < 0.001) and harvest year (χ² = 11.913, P = 0.003), with significant farming × year (χ² = 9.743, P = 0.008) and farming × climate × year interactions (χ² = 16.196, P < 0.001; Table S10). Across all seeds, conventional farming legacy and ambient climate legacy resulted in significantly higher germination than organic farming legacy and future climate legacy, respectively (Fig. S6A). Germination was also higher in 2024 than in 2018, whereas 2021 remained intermediate. Among the individual treatment combinations, the conventional– ambient legacies in 2024 showed the highest germination, whereas the organic–future combination in 2018 showed the lowest germination (Fig. S6B).

Before drought induction, plant height showed significant farming legacy main effects in 2018 (F = 9.963, P = 0.002) and 2024 (F = 4.875, P = 0.030), and significant climate legacy effects in 2021 (F = 4.053, P = 0.048) and 2024 (F = 3.996, P = 0.049; Table S11; Fig. 3B). In 2018, plants from the organic–future legacy were significantly shorter than those from both conventional legacy combinations (Fig. 3B). In 2024, plants from the conventional–ambient legacy were significantly taller than those from the organic–future legacy (Fig. 3B). After drought induction, water stress significantly reduced plant height in all harvest years. Across both watering conditions, farming legacy had a significant main effect on plant height in 2018 (F = 6.742, P = 0.011) and 2021 (F = 6.623, P = 0.012), whereas climate legacy had a significant main effect in 2024 (F = 4.471, P = 0.038; Table S11; Fig. 3B).

**Figure 3.**
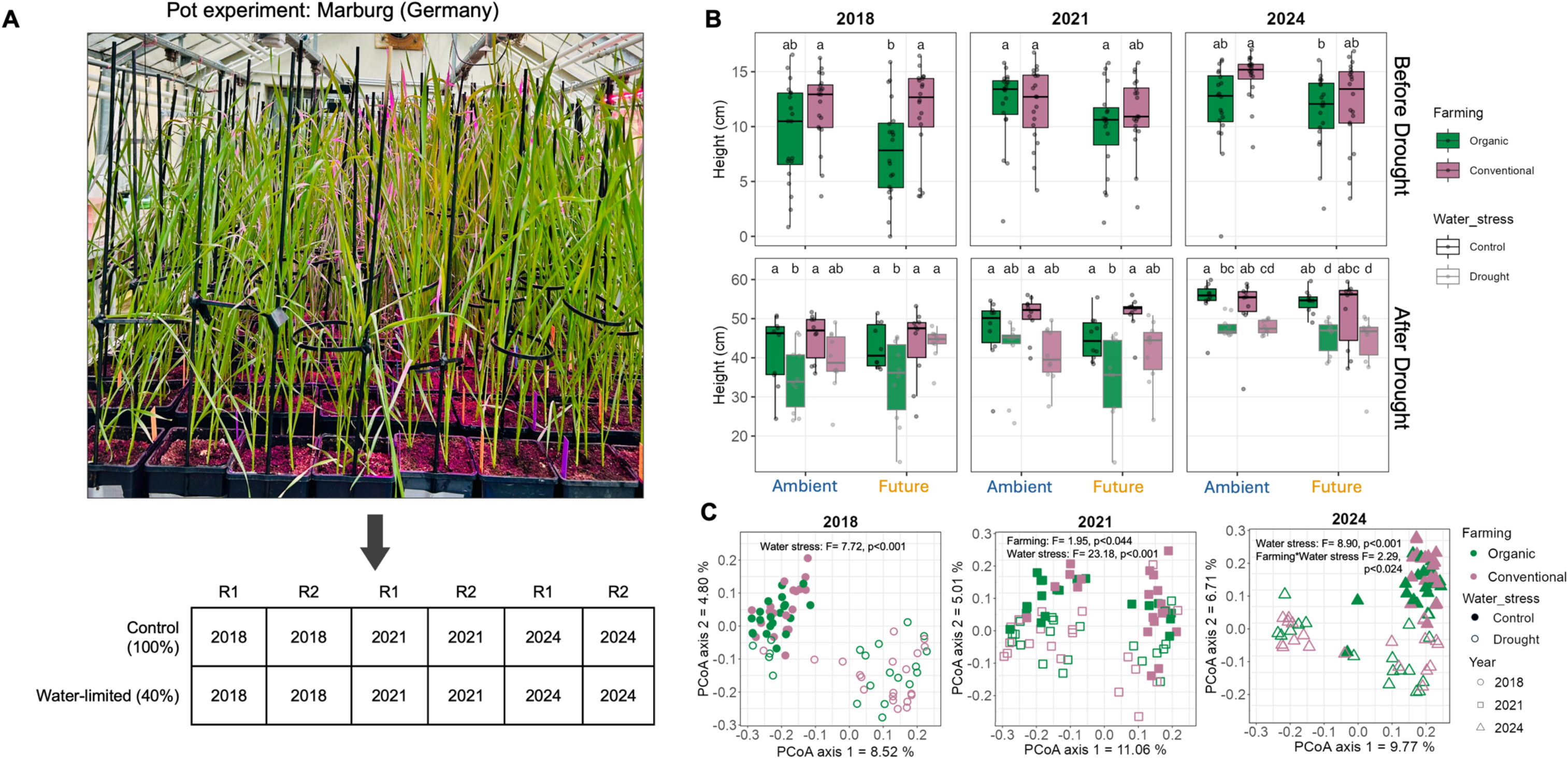
Offspring growth responses and rhizosphere bacterial community structure under water stress. (A) Greenhouse setup and experimental design. Winter wheat (*Triticum aestivum* L. cv. RGT Reform) seeds harvested in 2018, 2021 and 2024 from the GCEF were planted in non-sterile potting soil. Seeds represented organic and conventional farming legacies established under ambient and future climate treatments. Four seeds were sown per pot. At 14 d after sowing, pots were assigned to well-watered control or drought treatments, maintained at 100% and 40% of the control water supply, respectively, for six weeks. R1 and R2 denote the two greenhouse replicates. (B) Variations in plant height before and after drought exposure on plants originating from seed with distinct farming and climatic history which were harvested in 2018, 2021, and 2024. Different letters in the panel denote statistically significant differences based on post hoc Tukey’s test (*P*<0.05). Panels without letter indicate non-significant interaction. (C) Principal coordinate analysis (PCoA) based on Bray-Curtis dissimilarities of rhizosphere bacterial community composition. Samples are coloured by farming legacy of seeds (organic and conventional), filled symbols indicate well-watered controls and open symbols indicate drought-treated plants.

Aboveground fresh biomass was significantly reduced by drought across all harvest years (Table S12; Fig. S7A). In 2024, however, fresh biomass was also shaped by climate history (F = 8.921, P = 0.004), with a significant climate × water stress interaction (F = 5.544, P = 0.021, Table S12, Fig. S7A). By contrast, DWC increased significantly under drought in all years (Fig. S7B). In 2024, DWC was additionally affected by climate history (F = 10.11, P = 0.002), with overall higher values under ambient than future climate (Table S12, Fig. S7B). SPAD index was more year-specific than growth responses, with a farming × water stress interaction in 2018 and a climate × water stress interaction in 2024 (Table S13; Fig. S8A). Rhizosphere microbial CO_2_ production showed a year-specific drought response (Table S14; Fig. S8B). Water stress reduced CO_2_ production in 2018 (F = 10.83, P = 0.002) and 2024 (F = 13.27, P = 0.001), but not in 2021. In 2018, this response depended on farming legacy (Farming × Water Stress: F = 5.933, P = 0.018), whereas in 2024 the drought effect was consistent across farming histories.

#### 3.2.2. Diversity, community structure, and composition of the rhizosphere bacterial communities

Water stress significantly reduced offspring rhizosphere bacterial Shannon diversity in all harvest years (Table 1; Fig. S9A). Rhizosphere community structure was likewise shaped by water stress across harvest years (Table 1, Fig. 3C). Farming legacy contributed additionally in a year-specific manner, with a significant main effect in 2021 (F = 1.947, P = 0.044) and a significant farming × water stress interaction in 2024 (F = 2.293, P = 0.024, Table 1, Fig. 3C). The relative abundance of dominant rhizosphere taxa showed that harvest year and water stress were the primary determinants of compositional shifts, whereas farming and climate legacies influenced the abundance of specific taxa in rhizosphere (Table S15, Fig. S9B).

**Table 1.** ANOVA test results for the year wise-effects of Farming, Climate, Water Stress, and their interactions on the bacterial Shannon diversity and community structure of the rhizosphere. F = f-value; P = P-value.

| Factors | Shannon diversity |  |  | Community structure |  |  |
| --- | --- | --- | --- | --- | --- | --- |
|  | 2018 | 2021 | 2024 | 2018 | 2021 | 2024 |
| *Farming* | F = 3.695<br>P = 0.059 | F = 0.888<br>P = 0.349 | F = 2.334<br>P = 0.131 | F = 1.004<br>P = 0.386 | F = 1.947<br>**P = 0.044** | F = 1.416<br>P = 0.136 |
| *Climate* | F = 0.010<br>P = 0.919 | F = 0.078<br>P = 0.781 | F = 0.365<br>P = 0.548 | F = 0.598<br>P = 0.948 | F = 1.201<br>P = 0.225 | F = 0.767<br>P = 0.674 |
| *Water Stress* | F = 17.43<br>**P < 0.001** | F = 24.92<br>**P < 0.001** | F = 53.57<br>**P < 0.001** | F = 7.717<br>**P = 0.001** | F = 23.17<br>**P = 0.001** | F = 8.904<br>**P = 0.001** |
| *Farming × Climate* | F = 0.907<br>P = 0.344 | F = 0.496<br>P = 0.483 | F = 0.906<br>P = 0.345 | F = 0.670<br>P = 0.866 | F = 0.712<br>P = 0.752 | F = 0.776<br>P = 0.642 |
| *Farming × Water Stress* | F = 0.148<br>P = 0.702 | F = 1.149<br>P = 0.287 | F = 0.020<br>P = 0.889 | F = 1.027<br>P = 0.347 | F = 1.419<br>P = 0.135 | F = 2.293<br>**P = 0.024** |
| *Climate × Water Stress* | F = 0.752<br>P = 0.389 | F = 0.919<br>P = 0.341 | F = 0.045<br>P = 0.832 | F = 1.003<br>P = 0.383 | F = 0.792<br>P = 0.642 | F = 0.884<br>P = 0.497 |
| *Farming × Climate × Water Stress* | F = 0.558<br>P = 0.457 | F = 0.005<br>P = 0.941 | F = 0.007<br>P = 0.933 | F = 1.253<br>P = 0.168 | F = 0.960<br>P = 0.425 | F = 0.680<br>P = 0.769 |

\* Farming refers to the history of seed microbes from conventional and organic agriculture.
Climate refers to history of seed microbes from ambient and future climate under each farming.
Bold values indicate statistical significance (P < 0.05).

Rhizosphere bacterial communities showed substantial overlap across well-watered and drought conditions, with 2,705 ASVs shared between treatments across all harvest years (Fig. 4A). In addition, 205 ASVs were detected across all well-watered rhizosphere groups, whereas 138 ASVs were detected only in drought-treated rhizospheres. Among the drought-treated rhizospheres, the largest harvest-year-specific ASV fraction occurred in plants established from the 2018 seed batches (187 ASVs), followed by those established from the 2024 (177 ASVs) and 2021 seed batches (167 ASVs). Well-watered rhizospheres were dominated by *Legionella* and *Sphingobacteriaceae*, whereas the drought-specific fraction included *Micropepsaceae* and *Streptomyces* across all harvest years (Fig. 4A).

**Figure 4.**
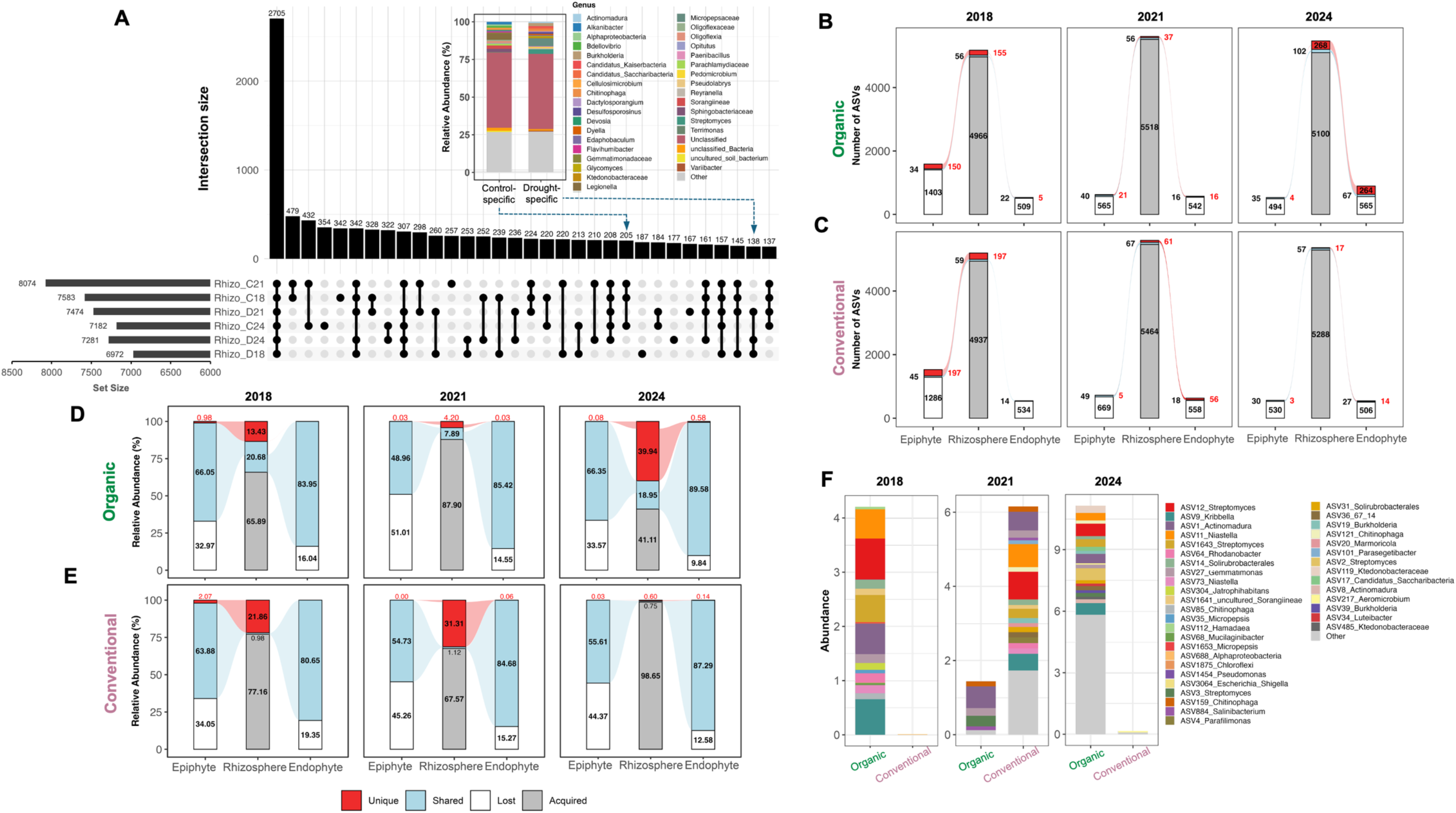
Rhizosphere bacterial responses to water stress and ASV overlap with seed communities across farming legacies. (A) Upset plot showing shared and unique bacterial ASVs within rhizosphere communities across the 3 harvest years (2018, 2021, and 2024) under well-watered control and drought conditions, where ‘C’ represent well-watered control conditions and ‘D’ represents drought conditions. Horizontal set size bars represent the total ASVs per group, filled circles indicate ASVs unique to a single compartment-year combination, dark circle connecting bar indicate shared ASVs between treatments. Inset highlights the relative abundance of ASVs (205) that are unique to control and drought ASVs (138) that are shared in all harvest years. (B, C) Sankey diagrams showing the numbers of ASVs assigned to the Shared, Unique, Lost and Acquired categories in epiphytic seed, endophytic seed and drought-treated offspring rhizosphere communities under organic (B) and conventional (C) farming legacies across the three harvest years. (D,E) Corresponding relative abundances of the four ASV categories under organic (D) and conventional (E) farming legacies. Numbers within bars indicate ASV counts in B,C and relative abundance (%) in D,E. Shared ASVs, shown in light blue, were detected in seeds and offspring rhizospheres across harvest years. Unique ASVs, shown in red, were harvest-year-specific ASVs detected in both seeds and the corresponding offspring rhizosphere. Lost ASVs, shown in white, were detected in seeds but not in the corresponding offspring rhizosphere. Acquired ASVs, shown in grey, were detected in the offspring rhizosphere but not in the corresponding seeds. Seed– rhizosphere ASV overlap comprises the combined Shared and Unique categories. (F) Abundance of top 20 bacterial ASVs detected in drought affected rhizosphere of offspring generation plants that originated from seed epiphytic and endophytic microbiota under organic and conventional farming legacies across harvest years.

#### 3.2.3. Shared ASVs between seeds and offspring rhizospheres under drought

Because water stress affected rhizosphere bacterial community structure and farming legacy contributed additional variation in specific harvest years (Table 1), we quantified ASV overlap between epiphytic and endophytic seed compartments and drought-treated offspring rhizospheres within each harvest year and farming legacy (Fig. 4B–E). Seed–rhizosphere ASV overlap (the combined shared and unique categories) accounted for a small fraction of rhizosphere richness. The largest overlap occurred under the 2024 organic farming legacy, with 370 ASVs (102 shared and 268 unique), equivalent to 6.76% of all rhizosphere ASVs. By comparison, 74 overlapping ASVs were detected under the 2024 conventional farming legacy, whereas 256 were detected under the 2018 conventional farming legacy (Fig. 4B, C). Harvest-year-specific unique ASVs occurred at low relative abundance in seeds, accounting for 0.00– 2.07% of epiphytic abundance and 0.00–0.58% of endophytic abundance, but represented up to 39.94% of rhizosphere relative abundance under the 2024 organic farming legacy and 31.31% under the 2021 conventional farming legacy (Fig. 4D, E). Shared ASVs accounted for a larger proportion of endophytic than epiphytic abundance across harvest years and farming legacies (80.65–89.58% versus 48.96–66.35%), whereas lost ASVs represented a smaller proportion (9.84–19.35% versus 32.97–51.01%). Under the 2024 conventional farming legacy, acquired ASVs comprised 5,288 ASVs and accounted for 98.65% of drought-rhizosphere relative abundance (Fig. 4B, E).

The abundance and taxonomic composition of shared ASVs detected in both seeds and drought-treated rhizospheres also differed among harvest years and farming legacies (Fig. 4F). In 2018 and 2024, organic farming legacies supported a broader and more abundant shared ASV fraction than conventional farming legacies. In 2018, major contributors included *Actinomadura* (ASV1) and *Streptomyces* (ASV12), whereas in 2024 the shared fraction was dominated by *Streptomyces* (ASV2 and ASV12), *Kribbella* (ASV9) and *Niastella* (ASV11). Under the conventional farming legacy, shared ASVs were detected at very low relative abundance in both years. The reverse pattern was observed in 2021, where shared ASVs were more abundant in the conventional than in the organic legacy rhizosphere (Fig. 4F). Year-specific unique ASVs were most numerous in 2018, with 16 abundantly shared between seeds and rhizosphere ASVs under conventional farming and 14 under organic farming, whereas only six and one such ASVs were detected in 2024 under organic and conventional farming, respectively, and none were detected in 2021 (Fig. S10).

#### 3.2.4. Association of plant traits with seed and rhizosphere bacterial diversity

Across legacy combinations and watering treatments, plant height showed the largest number of significant associations with seed and rhizosphere bacterial Shannon diversity (Fig. 5A). Under the organic–ambient legacy, epiphytic seed diversity was positively correlated with plant height in both well-watered (ρ = 0.675, P = 0.004) and drought-treated plants (ρ = 0.720, P = 0.003). In well-watered plants from the same legacy combination, plant height was also positively correlated with endophytic seed diversity (ρ = 0.521, P = 0.042) and rhizosphere diversity (ρ = 0.573, P = 0.023). Under the conventional–ambient legacy, rhizosphere diversity was positively correlated with plant height under drought (ρ = 0.648, P = 0.007). In the same treatment, endophytic seed diversity was negatively correlated with fresh biomass (ρ = −0.573, P = 0.023), whereas grain dry weight was positively correlated with dry weight content (ρ = 0.542, P = 0.033) and negatively correlated with fresh biomass (ρ = −0.510, P = 0.048; Fig. 5A). Epiphytic seed Shannon diversity was also positively associated with offspring rhizosphere Shannon diversity. When grouped by farming legacy, this relationship was significant under organic farming (ρ = 0.253, P = 0.006) but not under conventional farming (ρ = 0.145, P = 0.115). On the other hand, when grouped by climate legacy, significant positive relationships were detected under both ambient (ρ = 0.190, P = 0.039) and future conditions (ρ = 0.216, P = 0.019; Fig. 5B). We next asked whether abundances of shared seed-rhizosphere taxa in the drought rhizosphere were associated with offspring performance. Plant height showed the largest number of significant genus-level correlations, with more significant associations detected under organic than conventional farming legacy (Fig. 5C). By contrast, fresh biomass and dry weight content were associated with fewer taxa, with dry weight content under organic legacy showing negative correlations with genera including *Chryseolinea*, *Ohtaekwangia*, *Dactylosporangium* and *Saccharimonadales* (Fig. 5D, E).

**Figure 5.**
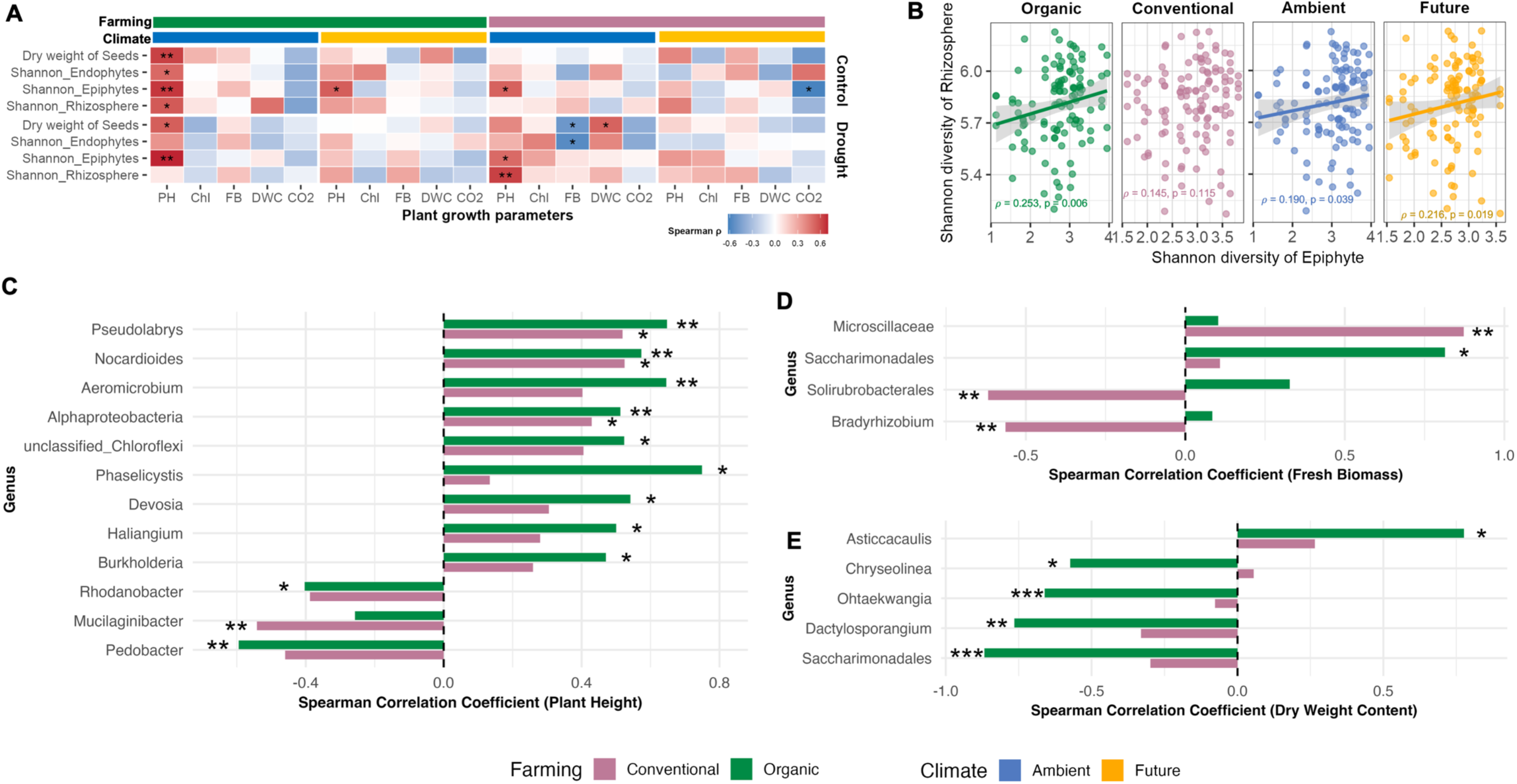
Correlations between seed bacterial diversity, rhizosphere diversity, and plant performance under drought across farming and climate legacies. (A) Heatmap showing Spearman correlation coefficients (with Benjamini–Hochberg correction) between plant growth and physiological traits (plant height: PH, chlorophyll content: Chl, fresh biomass: FB, dry weight content: DWC, and CO_2_ respiration: CO2) and Shannon diversity of seed epiphytic, endophytic, and drought-affected rhizosphere bacterial communities across farming and climate histories. Colour intensity represent the strength and direction or correlations, Negative correlation (Blue), Positive correlation (Red). (B) Scatter plots with fitted linear regression lines illustrating significant correlations between seed and rhizosphere Shannon diversity under different farming (organic vs. conventional) and climate (ambient vs future) legacies based on Spearman correlation analysis with Benjamini–Hochberg correction. Spearman correlation coefficient (ρ) and corresponding p-values are shown within each panel. (C-E) Differential correlations between transferred seed-associated bacterial ASVs and plant growth traits under drought. Bar plots represent significant Spearman correlations with Benjamini–Hochberg correction (p < 0.05) between bacterial ASV abundance in the drought affected rhizosphere and (C) plant height, (D) fresh biomass, and (E) dry weight content. The color of bars indicates the farming legacy of the seed source: organic farming legacy (green), conventional farming legacy (violet). Asterisks represent significance levels (*: p < 0.05, **: p < 0.01, ***: p < 0.001 respectively).

## 4. Discussion

Using winter wheat seeds harvested across a multi-year field experiment, our study shows that seed-associated bacterial communities retained distinct signatures of farming and climate legacy established in the maternal generation. These legacy patterns were also associated with offspring establishment, growth, and rhizosphere bacterial communities under drought stress. Harvest year, reflecting contrasting climatic histories, was the dominant driver of seed traits and overall seed bacterial community structure. By comparison, farming legacy was most clearly reflected in endophytic bacterial diversity and was associated with several offspring performance traits. Collectively, our findings revealed that seeds retain bacterial signatures of parental field history that are associated with offspring rhizosphere composition and plant performance.

Harvest year imposed the strongest imprint on both epiphytic and endophytic communities. Because harvest year covaried with post-harvest storage duration, the observed differences likely integrate climatic conditions during seed development with storage-associated changes in seed bacterial communities [22,48,49]. Prolonged storage can reduce bacterial diversity and alter community composition by favoring taxa that tolerate storage conditions, with seed-surface communities potentially being particularly responsive [48,50]. Among the sampled years, 2018 represented the strongest climatic perturbation, with warmer and drier growing-season conditions than the 2016–2017 baseline, consistent with the broader drought–heat event reported across Europe [4,5]. In line with this, grain dry weight and seed bacterial diversity were negatively associated with temperature and positively associated with precipitation, showing that climatic conditions during seed development influenced both seed traits and microbiome assembly. This pattern is biologically plausible because drought during grain filling constrains assimilate allocation and alters the physical and nutritional environment available for microbial colonization within developing seeds [51,52]. A subset of ASVs detected in 2018 seeds was also detected in rhizospheres established from the corresponding seed batches under drought, indicating that a harvest-year-associated bacterial signature remained detectable in the offspring rhizosphere. However, this co-detection does not establish direct seed-to-rhizosphere transmission. ASVs assigned to bacterial groups including *Pantoea, Massilia*, *Pseudomonas*, and *Sphingomonas* contributed to the taxonomic differences among harvest years. These genera are commonly known for their stress tolerance and survival under water-limited conditions [53–56]. For instance, *Sphingomonas* and related *Alphaproteobacteria* are frequently associated with phyllosphere and seed environments due to their tolerance to desiccation and UV stress, as well as their ability to utilize plant-derived carbon compounds [57]. However, their functional relevance to offspring performance requires direct experimental validation.

Among parental field treatments, farming legacy was the most important determinant of endophytic bacterial diversity, whereas climate legacy was more evident in epiphytic diversity and community structure. Epiphytic communities responded to both harvest year and climate legacy, whereas endophytic communities were comparatively less responsive to these factors. This contrast is consistent with the greater environmental exposure of the seed surface and stronger host filtering within seed tissues during seed development [31,58,59]. Environmental exposure and host-mediated recruitment may therefore leave different bacterial signatures in the two seed compartments. A notable result was that the conventional farming legacy was associated with significantly higher endophytic bacterial diversity in both 2018 and 2021, despite the common expectation that organic systems sustain greater microbial diversity in bulk soil [44,60]. This contrast suggests that the diversity of the surrounding soil community does not necessarily predict the subset of taxa that successfully enter developing seeds. Instead, seed microbiome structure appears to depend on strong compartment-specific filtering, with farming legacy being expressed more in the endophytic compartment. The seed–rhizosphere analysis also reflected this compartmental contrast, where shared ASVs accounted for a larger proportion of endophytic than epiphytic relative abundance, whereas lost ASVs accounted for a smaller proportion. This pattern shows that a larger proportion of the endophytic community was represented by ASVs also detected in offspring rhizospheres. Consistent with this interpretation, endophytic diversity under ambient climate showed a stronger positive association with grain dry weight than epiphytic diversity, further suggesting that bacteria residing within seed tissues are more closely linked to seed trait than those restricted to the seed surface [54,61].

Farming legacy and climate legacy established in the maternal generation were associated with offspring establishment and growth, although the direction and strength of these associations differed among legacy combinations and harvest years. Across the complete dataset, germination was higher for seeds from the conventional than the organic farming legacy and from the ambient than the future climate legacy. This pattern is consistent with our previous work at the same GCEF site, where soil microbiomes shaped by conventional farming under ambient climate most strongly promoted wheat establishment under drought [47,62]. However, this should not be interpreted as the general superiority of the conventional farming legacy. Conventional farming legacy was associated with greater plant height in several harvest-year-specific comparisons, whereas plant height showed a larger number of significant taxon-level associations under organic farming legacy. Together, these results show that farming legacy and climate legacy were associated with early establishment and with context-dependent relationships between legacy-associated bacterial patterns and plant growth under drought.

Across all harvest years, water stress reduced rhizosphere bacterial Shannon diversity and altered community structure, whereas farming legacy contributed additional variation in specific harvest years and climate legacy had no detectable effect on rhizosphere community structure. Nevertheless, bacterial signatures associated with the maternal seed source were not fully obscured. ASV overlap between seeds and drought-treated offspring rhizospheres differed among harvest years and farming legacies, and harvest-year-specific unique ASVs collectively accounted for little relative abundance in seeds but represented up to 39.94% of rhizosphere relative abundance under drought. This pattern is consistent with ecological amplification during rhizosphere establishment, but it does not demonstrate that individual ASVs were transmitted directly from seeds or caused changes in plant performance. Recent controlled experiments in wheat showed that seed-borne bacteria can contribute substantially to early rhizosphere microbiome assembly [63]. In parallel, direct intergenerational transmission of *Pantoea agglomerans* was demonstrated by recovering nearly identical isolates across three wheat generations and experimentally tracking a labelled isolate into the subsequent generation [64]. Comparable controlled experiments that track genetically distinguishable bacterial strains from maternal plants or seeds into offspring tissues and rhizospheres are required to validate the ASV-overlap patterns identified here [19,65]. Such evidence would establish whether candidate seed-associated bacteria persist and function under drought and would provide a stronger basis for their subsequent evaluation in synthetic communities or microbiome-informed seed treatments. As compound drought–heat events become more frequent, understanding and harnessing the microbial legacy carried by seeds may offer an underused but ecologically grounded route to improving agroecosystem resilience.

## 5. Conclusion

This study shows that seeds retain bacterial signatures of parental climate history and farming legacies, and that these signatures remain biologically relevant in the next generation under drought. Harvest year, encompassing contrasting climatic histories and different storage durations, was the dominant driver of seed traits and overall seed bacterial community structure, whereas farming legacy was most clearly reflected in the endophytic compartment. Moreover, some harvest-year-specific unique ASVs occurred at low relative abundance in seeds but reached substantially higher relative abundance in drought-treated offspring rhizospheres. In the offspring generation, water stress was the dominant filter on rhizosphere diversity and community structure, while farming legacy contributed additional year-specific variation and legacy-associated bacterial patterns were linked to several plant traits. Together, these findings identify seeds as structured microbial habitats that retain signatures of maternal field history and are associated with offspring rhizosphere composition and plant performance.

## Data availability

The data that support the findings of this study are available in the supplementary material of this article. Fastq files are deposited in the NCBI Sequence Read Archive (the BioProject accession PRJNA1508365).

## Supporting information

Supplementary results

Supplementary datasheet

## Acknowledgments

We would like to thank the research group at the Global Change Experimental Facility for providing us with the seed samples and access to the field. This work was funded by the Deutsche Forschungsgemeinschaft (DFG, German Research Foundation; project number: 523864000 to HA and 446841743 to JGR) and the Max Planck Society (JGR and NNO).

## Author contributions

HA designed the study. BS planned and conducted the greenhouse experiment, performed the bioinformatic analyses, analyzed the data, and wrote the manuscript with the help of HA. JB helped with DNA extraction and the greenhouse experiment. MS provided the seed and access to the GCEF field. JGR and NNO helped with CO_2_ production measurements. All co-authors participated in reviewing and editing the final text. The authors declare no competing interests.

