## Supplementary methods for "Farming and climate legacies shape the seed microbiota and offspring drought responses in wheat"

### Supplementary material and methods

#### Extraction of seed epiphyte, endophyte and rhizosphere microbiota

DNA from seed epiphytic and endophytic bacterial communities was extracted following a modified protocol described by Azarbad et al. (2022). For epiphytic bacteria, 1 g of seeds per plot was incubated in 5 mL sterile peptone buffer [2] in 50 mL Falcon tubes and shaken at 500 rpm for 1 h. The suspension was centrifuged at 8,000 × g for 15 min, the supernatant was discarded, and the pellet was resuspended in 800 µL peptone buffer and stored at −20 °C until DNA extraction. For endophytic bacteria, seeds remaining after epiphyte removal were surface-sterilized by sequential washing with sterile water, immersion in 95% ethanol for 5 min, sodium hypochlorite solution (2.5% available $Cl^-$) for 5 min, and 95% ethanol for 3 min, followed by three rinses with sterile water. Sterilization efficiency was confirmed by plating 100 µL of the final rinse on R2A agar and incubating at 28 °C for 3 days; no microbial growth was observed (Fig. S1). Sterilized seeds were immediately flash-frozen in liquid nitrogen, ground to a fine powder using a mortar and pestle, and 0.5 g of homogenized material was stored at −20 °C prior to DNA extraction. For DNA extraction, we used DNeasy® PowerSoil® Pro Kit (QIAGEN) based on the manufacturer's protocols. For epiphyte bacteria, pellet resuspended in 800 µL peptone buffer was directly used, while for endophytes, 1 g of fine powder of seeds after sterilization was used. Rhizosphere microbial communities were characterized using 250 mg of rhizosphere soil following the same DNA extraction described for seed epiphytic and endophytic microbiota.

#### Rhizosphere CO2 production

Microbial activity in the rhizosphere was quantified via *in-vivo* carbon dioxide ($CO_2$) production. For that, we used the collected rhizosphere soil (0.75 g), which was stored in glass vials, then flushed with Argon to remove any background gases and incubated for 48 h for microbial activation and $CO_2$ production. The produced $CO_2$ amounts *in-vivo* were determined via headspace analysis using a Clarus 690 GC system [gas chromatography–flame ionization detector/thermal conductivity detector (GC-FID/TCD); PerkinElmer Inc., Waltham, USA] with a custom-made column circuit (ARNL6743) that was operated with the TotalChrom v.6.3.4 software (PerkinElmer Inc., Waltham, USA). The headspace samples were injected by a TurboMatrixX110 (PerkinElmer Inc., Waltham, USA) autosampler, heating the samples to 65°C for 20 min before injection. The samples were then separated on a HayeSep column (7′ HayeSep N 1/8″ Sf; PerkinElmer Inc., Waltham, USA) kept at 60°C. Subsequently, the gases were detected with a TCD (at 200°C). The quantification of all substrates was based on a linear standard curve derived from measuring varying amounts of $CO_2$ under identical conditions [3].

#### Amplicon sequencing and data processing

DNA samples (seed epiphyte: 60, seed endophyte: 60, rhizosphere: 240) resulting from both experiments were sent to Biomarker Technologies (Germany) for libraries preparation and Illumina MiSeq (paired-end) sequencing. The V5–V6 region was amplified using primers 799F (AACMGGATTAGATACCCKG) and 1115R (AGGGTTGCGCTCGTTG). Processing sequence data was conducted in QIIME 2 to generate Amplicon Sequence Variants (ASVs). Briefly, primer sequences were removed, and reads were truncated ensuring that only high-quality bases remained prior to denoising. Chimeric sequences were identified and removed with the removeBimeraDenovo function of DADA2. For taxonomic affiliations of the resulting amplicon sequence variants (ASV), a naive Bayesian classifier was performed based on the SILVA database v138. ASVs unassigned at the phylum level, together with chloroplast and mitochondrial reads, were removed from the dataset. Before further analysis, the following filtering criteria were applied: samples should have more than 1000 ASV reads, and any ASVs with less than five reads in a given sample were removed [4]. Furthermore, any ASV that was found in only one sample was discarded from the data set. For the calculation of alpha diversity, the data were normalized on the basis of sequencing depth. For beta diversity, we normalized the data set based on the relative abundance of ASVs in each sample [1, 5].

**2.3. Statistical analysis**

All statistical analyses were conducted in R (Version 4.3.2; R Development Core Team, 2024) using R Studio (Version 2023.12.1+402). Climate variables at GCEF (air temperature and precipitation) were analyzed using linear-mixed models (LMMs) implemented in the *lme4* package. Harvest year, month, and their interactions were included as fixed effects, while field subplot was included as a random intercept. Statistical significance of the fixed effects was assessed using *lmerTest* package, applying type III ANOVA tests on the models. Unless otherwise stated, response variables measured across the three harvest years (e.g., grain dry weight and α-diversities) were first analysed using ANOVA that included harvest year and the relevant experimental factors and their interactions. Separate models were then fitted within each harvest year to characterize year-specific effects of farming and climate legacies and, for the greenhouse experiment, watering treatment. Within-year analyses compared seed batches stored for the same duration and therefore allowed farming legacy and climate legacy effects to be evaluated without direct confounding by differences in storage duration among years. Following ANOVAs, significant treatment effects were assessed by Tukey’s HSD test for pairwise comparisons. The α-diversities of the seed-associated microbiota (epiphytes and endophytes) were calculated using the Shannon index [6] via *agricolae* package [7]. Spearman's rank correlation was applied to investigate the correlation between the dry weight of seeds and Shannon diversity of seed-associated bacteria. The p-values were adjusted for multiple testing using the Benjamini-Hochberg procedure (FDR < 0.05). Next, we conducted Principal Coordinate Analyses (PCoA) based on Bray–Curtis dissimilarity using the *vegan* package [8] to visualize microbial community structure and the Permanova test (through the '*adonis2*' function) to evaluate the effect of experimental factors on the bacterial community structure. A total of 2 extreme outlier samples from 2024 were removed prior to visualization to improve ordination clarity and avoid compression of the main sample distribution in the PCoA plots. The relative abundance of the dominant top 30 bacterial ASVs were analyzed using ANOVA, considering the main and interaction effects of experimental factors. Furthermore, year-wise unique and shared epiphytic and endophytic ASVs were plotted as UpSetPlot using *ComplexUpset* package [9, 10].

As for the greenhouse experiment, germination data were analysed using a binomial generalized linear mixed model (GLMM) with farming legacy, climate legacy, and harvest year as fixed factors, followed by Tukey-adjusted pairwise comparison of estimated marginal means. Plant height, fresh biomass, DWC, chlorophyll content (SPAD index), and rhizosphere respiration ($CO_2$) were analysed using factorial ANOVA models to evaluate variations in plant performance across experimental factors, including farming legacy, climate legacy, harvest year, and water stress, followed by Tukey's HSD tests. Rhizosphere bacterial diversity, structure, and composition were analyzed using a similar workflow as used for seed data, with water stress included as an additional factor. The abundance of the top 50 dominant rhizosphere bacterial taxa (mean abundance of >1%) was studied at the genus taxonomic level. Stacked bar plots were generated to visualize the variation in the relative abundance of the dominant bacterial taxa that showed changes due to experimental factors. ASVs unique to or shared among rhizosphere communities under well-watered and drought conditions were summarized separately for each harvest year using UpSetPlot plots generated with the *ComplexUpset* package [9, 10]. To assess bacterial ASV overlap between seeds and offspring rhizospheres, ASV occurrence and relative abundance were compared separately between epiphytic or endophytic seed bacterial communities and drought-treated rhizospheres within each harvest year and farming legacy. The resulting detection patterns were visualized using alluvial diagrams generated with the *ggalluvial* package [11]. For these analyses, ASVs were classified into four categories: (i) Shared, representing ASVs detected in both seed and offspring rhizosphere across harvest years; (ii) Unique, representing harvest year-specific ASVs detected in both seeds and the corresponding offspring rhizosphere; (iii) Lost, representing seed-associated ASVs that were not detected in the offspring rhizosphere; and (iv) Acquired, representing ASVs detected in the offspring rhizosphere but not in the corresponding seed samples. Finally, relationships among seed and rhizosphere microbial diversity, plant growth traits, and grain characteristics (derived from the field experiment) were assessed using Spearman's correlation ($p < 0.05$, Benjamini-Hochberg corrected). Significant correlations ($p < 0.05$) were visualized as heatmaps created using *ComplexHeatmap* package [12]. All graphical outputs were generated using *ggplot2* [13], unless otherwise stated.
