## Supplementary results for "Farming and climate legacies shape the seed microbiota and offspring drought responses in wheat"

### Extended results

#### Seed bacterial community structure and composition

We examined shifts in the taxonomic composition of seed-associated bacterial communities for both epiphytic and endophytic fractions (Table S8-9; Fig S4). Across all samples, *Alphaproteobacteria* (ASV688), *Pantoea* (ASV3822, ASV5417), *Massilia* (ASV3327), and *Curtobacterium* (ASV12103) represented the most abundant taxa in the seed epiphyte (Fig S3A). The relative abundance of most genera varied significantly across harvest years (all F values ranging from 174.97 to 3.98, $P < 0.05$; Table S8). In comparison, farming effects were less pronounced, although certain taxa responded significantly to long-term management history. For instance, *Massilia* (ASV5358: Direct effect of Farming, $F = 4.120$, $P = 0.048$) showed the highest abundance under organic farming, while *Pseudomonas* (ASV117932: Farming × Year, $F = 4.723$, $P = 0.013$) displayed enrichment under conventional farming in 2024 (Table S8, Fig S4A). Climatic conditions further structured the epiphytic communities, with several genera significantly enriched under ambient climate treatments. For example, *Hymenobacter* (ASV12220: $F = 3.607$, $P = 0.035$), *Sphingomonas* (ASV10268: $F = 7.160$, $P = 0.002$; ASV116264: $F = 14.973$, $P<0.001$), *Massilia* (ASV5358: $F = 6.047$, $P = 0.005$), *Pantoea* (ASV5417: $F = 4.476$, $P = 0.017$) were particularly abundant in 2021, whereas *Hymenobacter* (ASV5851: $F = 4.278$, $P = 0.020$) and *Pedobacter* (ASV60528: $F = 6.077$, $P = 0.004$) exhibited higher relative abundance in 2024 (Climate × Year; Table S8, Fig S4A). In addition, the interaction of farming practice, climate history, and harvest year significantly shaped epiphytic community composition. For instance, in 2018, under conventional farming, *Pseudomonas* (ASV1454: $F = 4.040$, $P = 0.024$) showed increased abundance under ambient climate conditions, while declined under future climate conditions (Farming × Climate × Year; Table S8, Fig S4A).

In contrast to epiphytic communities, seed endophytes exhibited comparatively greater compositional stability, with fewer ASVs responding significantly to farming and climate interactions across years (Table S9). Nevertheless, harvest year remained a significant driver of variation in the relative abundance of most of the detected ASVs (ANOVA, all $P < 0.05$, F-value ranged from 3.205 to 19.437; Table S9, Fig S4B). Several taxa, including *Curtobacterium* (ASV12103: $F = 7.743$, $P = 0.001$) and *Pantoea* (ASV3822: $F = 5.908$, $P = 0.005$; ASV5417: $F = 8.996$, $P<0.001$) were absent in 2018 but increased in subsequent years, reaching its highest relative abundance in 2024 (Table S9, Fig S4B). A significant direct effect of Farming and harvest year was observed on the relative abundance of *Alphaproteobacteria* (ASV116307, ASV116407) and *Rhizobiales* (ASV116340, ASV116349). *Alphaproteobacteria* (ASV116307) also responded significantly to interactions between harvest year and climate conditions ($F = 3.476$; $P = 0.039$), with its relative abundance highest in seeds from future climate conditions in 2018 (Table S9, Fig S4B).

Additionally, grain dry weight was associated with distinct seed-associated bacterial taxa across years and legacy conditions (Fig. S5). In 2018, positive correlations under conventional farming included *Actinobacteria* and *Acidothermus,* whereas under ambient climate they included *Paenibacillaceae, Propionicimonas,* and *Duganella* (Fig. S5). In 2024, when grain dry weight was highest in the conventional treatments (Fig. 1C), positive correlations under conventional farming included *Frigoribacterium, Gracillibacteria,* and *Polyangiales*. Climate-specific analysis for 2024 further identified positive correlation with *Rhizobiales* under ambient climate and *Solirubrobacter, Salinispora,* and *Nitrosomonadaceae* under future climate in 2024 (Fig. S5).

**Rhizosphere bacterial community structure and composition**

We further examined changes in bacterial community composition at the genus level in rhizosphere samples under experimental conditions by analyzing the 50 most abundant taxa. Overall, harvest year and water stress were the primary determinants of genus-level shift in abundance explaining most of the observed compositional changes (Table S15, Fig S9B). *Streptomyces* was the dominant genus, particularly enriched in 2018 (Direct effect of Year, F = 28.858, P<0.001), however, it exhibited strong drought sensitivity, with its relative abundance significantly reduced under water stress (Year × Water stress, F = 6.318, P = 0.013). Farming legacy exerted selective and condition-dependent effects on several taxa. For instance, *Actinomadura* showed a clear direct response to farming history (F = 5.370, P = 0.021), with higher relative abundance under history of organic farming with well-watered condition in 2018 (Year × Water stress, F = 16.8413, P < 0.001), Similarly, with the history organic farming, *Niastella* (F = 7.053, P = 0.008) was more abundant under well-watered control treatment, however, relative abundance of *Jatrophihabitans* (F = 3.966, P = 0.048) decreased under this treatment combination (Farming × Water stress, Table S15, Fig S9B). In contrast, *Alphaproteobacteria* (F = 4.420, P = 0.037), *Gemmatimonadaceae* (F = 8.720, P = 0.003), *Iamia* (F = 7.931, P = 0.005), *Nocardiodes* (F = 5.386, P = 0.021), and *Rayranella* (F = 6.134, P = 0.014) showed a positive shift toward conventional-control treatment (Farming × Water stress, P<0.05). Climate legacy effects were comparatively weaker but still evident for specific taxa. For example, *Bradyrhizobium* was consistently more abundant in plants originating from future climate conditions (direct effect of Climate, F = 5.828, P = 0.017). In 2018, *Burkholderia* further increased in relative abundance in ambient compared to future climate histories (Climate × Year, F = 4.262, P = 0.040; Table S15, Fig S9B). A similar ambient-climate driven enrichment under conventional farming was observed for *Eaphobaculum* in well-watered control treatments (Farming × Climate × Water stress, F = 3.896, P = 0.050). Moreover, a four-way interaction of Farming and Climate history with harvest year was evident in drought-exposed next-generation plants for *Pseudorhodoplanes* (F = 4.888, P = 0.028), where conventional farming under future climate conditions promoted *Pseudorhodoplanes* under well-watered control conditions (Farming × Climate × Year × Water stress, Table S15, Fig S9B).

**Figures**

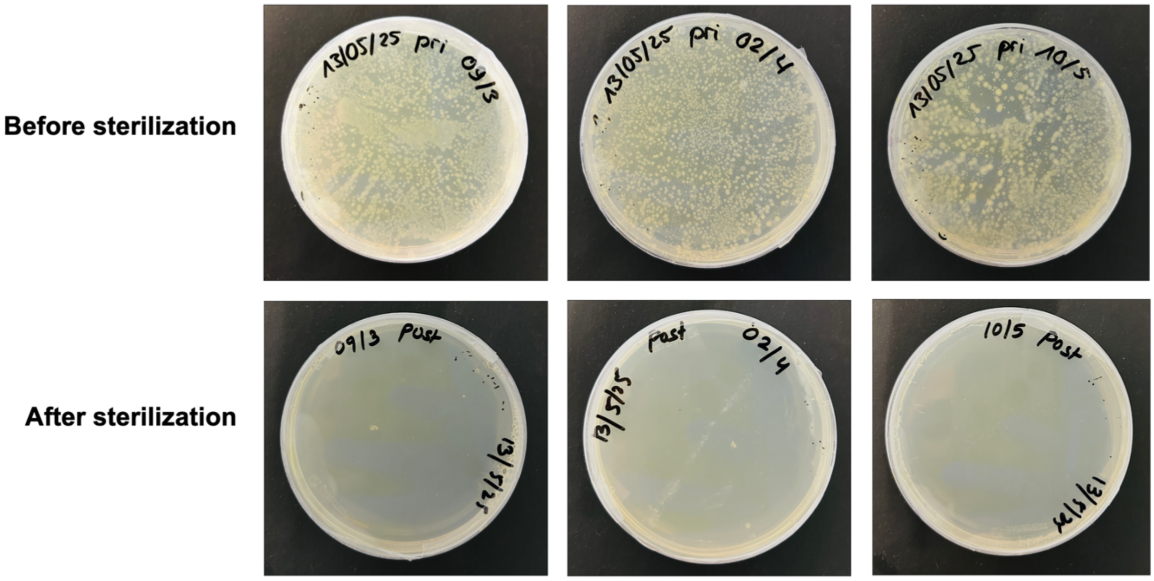


Figure S1: Photographic images of bacterial growth validation during seed microbiome extraction. Plates show bacterial colonies recovered (A) before surface sterilization for characterization of seed epiphytic microbiome and (B) absence of bacterial growth after seed surface sterilization prior to endophytic microbiome extraction.

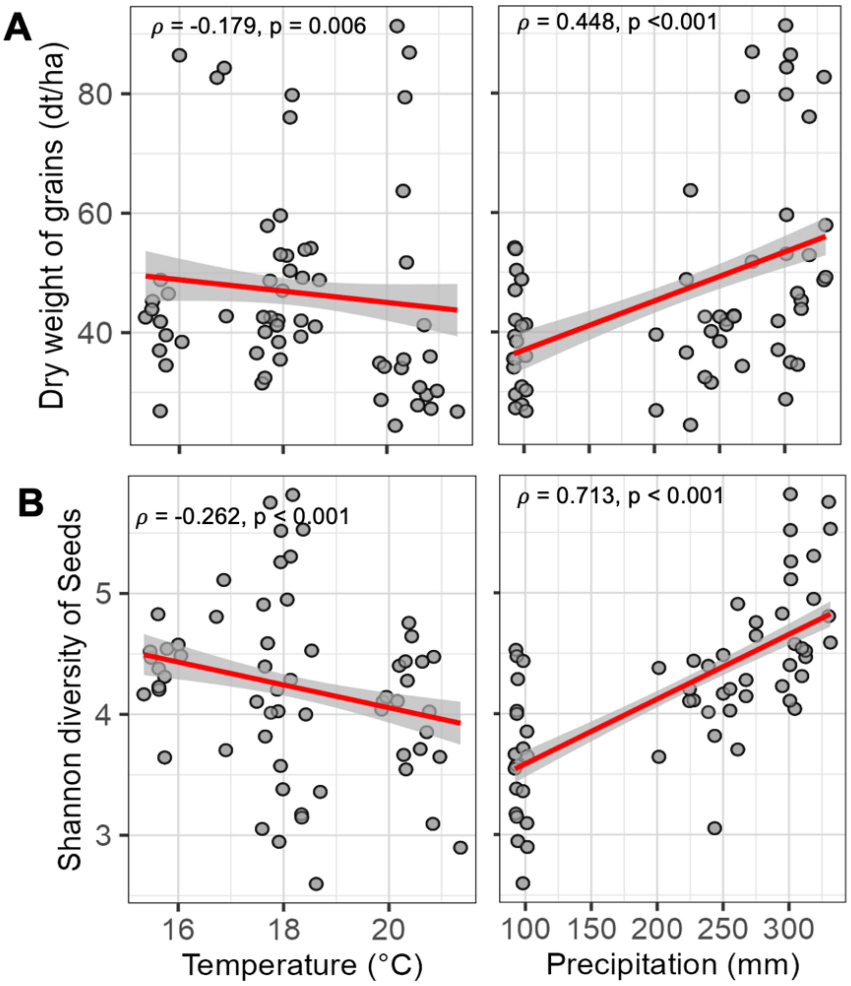


Fig S2: Scatter plots with fitted linear regression lines illustrating significant correlations between (A) dry weight of grains and (B) Shannon diversity of seed microbiota with temperature and precipitation of respective years based on Spearman correlation analysis with Benjamini–Hochberg correction. Spearman correlation coefficient (ρ) and corresponding p-values are shown within each panel.

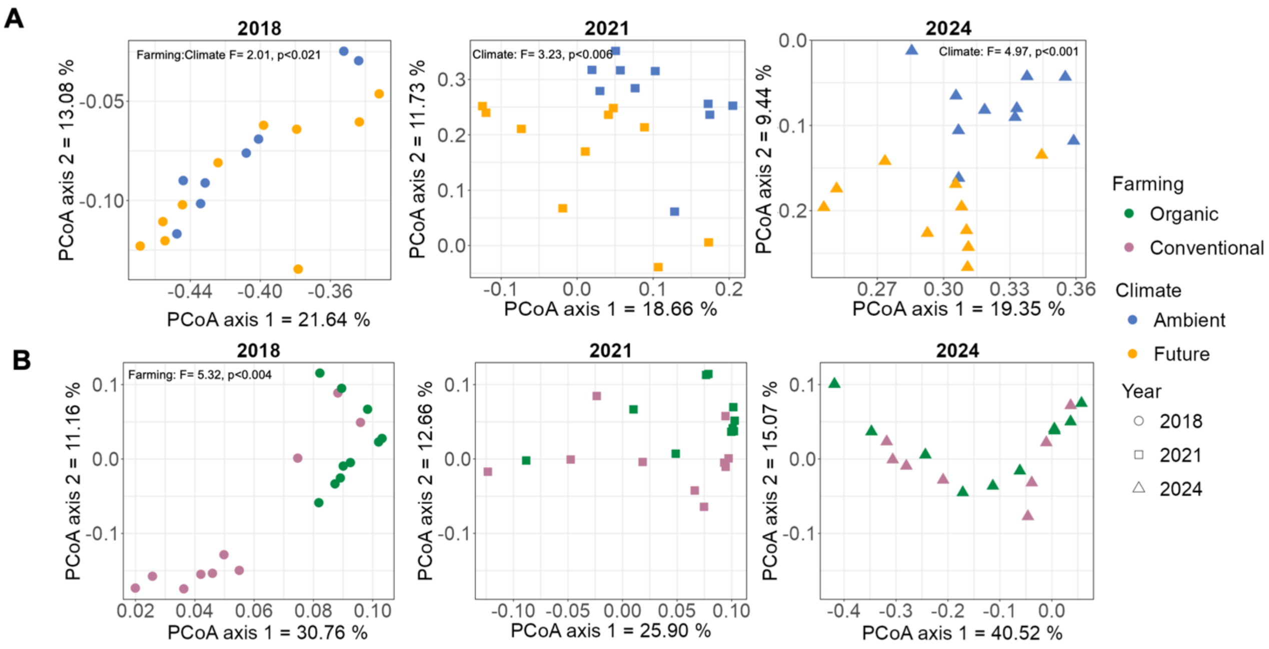


Figure S3: Principal coordinate analysis (PCoA) based on Bray-Curtis dissimilarities of (A) epiphytic and (B) endophytic bacterial ASV composition. Samples are coloured by coloured by climate legacy for epiphytic communities and farming legacy for endophytic communities and point shapes indicate harvest year.

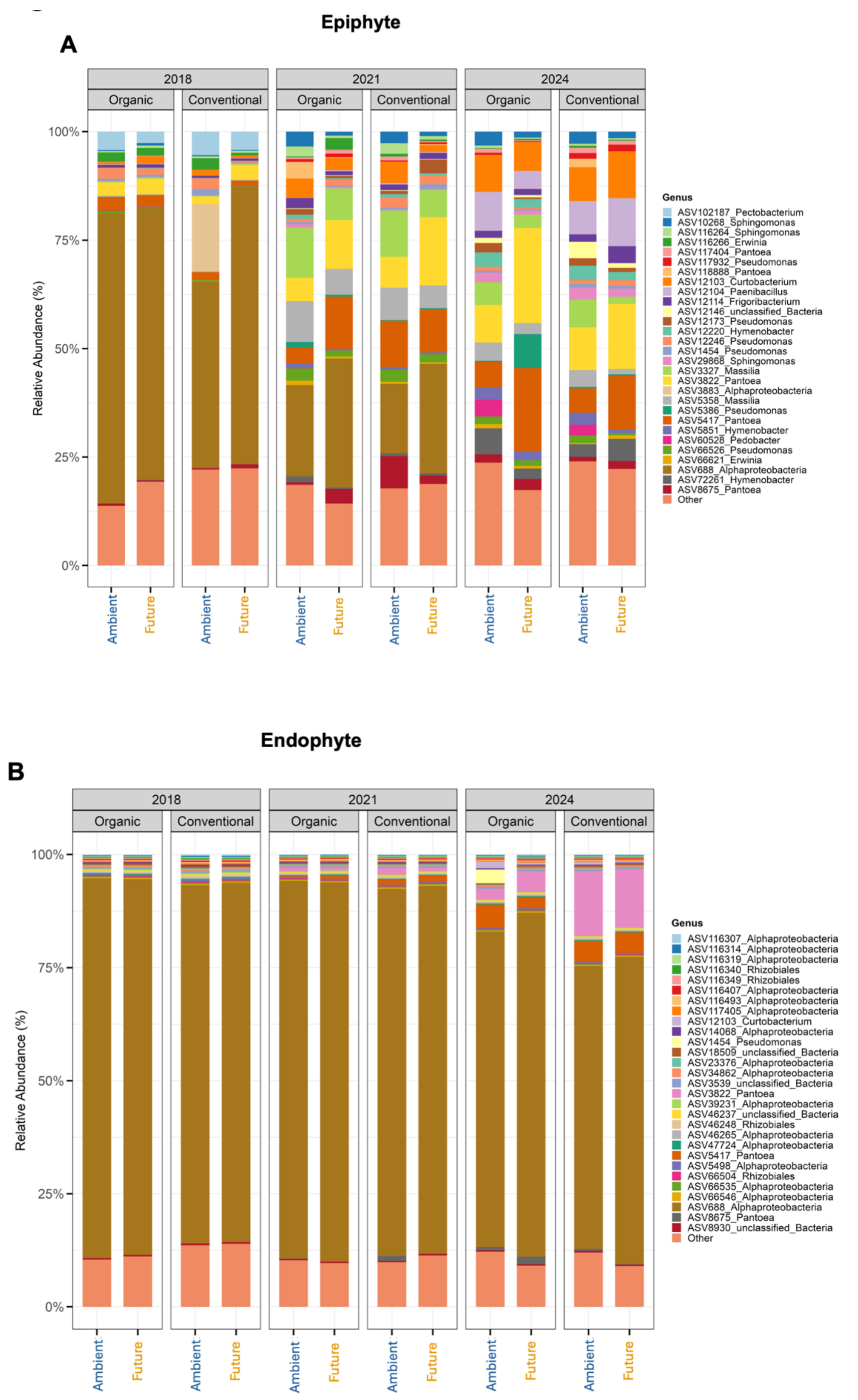


Figure S4: Relative abundance of top 30 most dominant bacterial ASVs detected in seed (A) epiphytic and (B) endophytic microbiome.

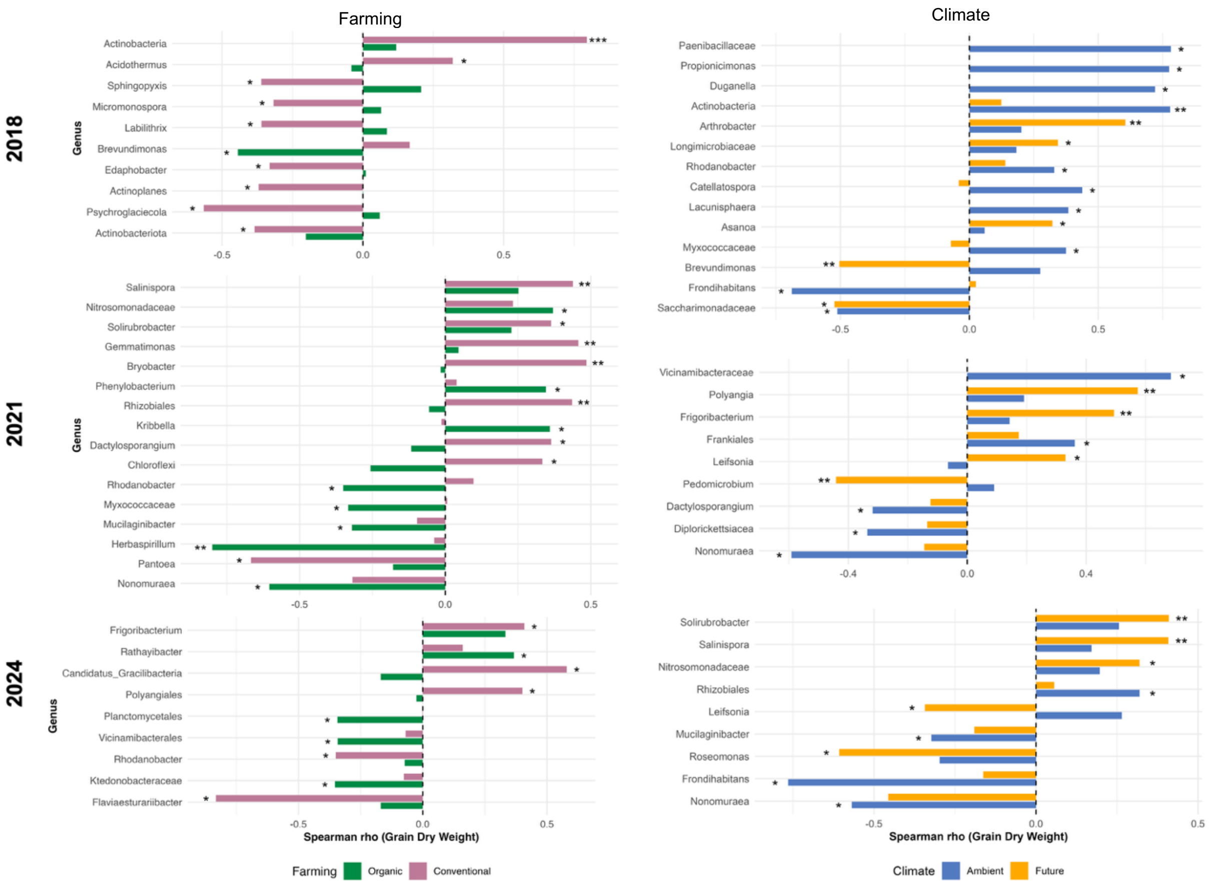


Figure S5: Differential correlations between seed-associated bacterial genera and dry weight of grains harvested in 2018, 2021, and 2024 under different farming and climate conditions. Bar plots represent significant Spearman correlations ($p < 0.05$) between bacterial genera abundance in the seed and dry weight of seeds. The color of bars indicates the farming conditions: organic farming (green), conventional farming (violet) and climate conditions: ambient (blue), future (yellow) of experimental field. Asterisks represent significance levels (*: $p < 0.05$, **: $p < 0.01$, ***: $p < 0.001$ respectively).

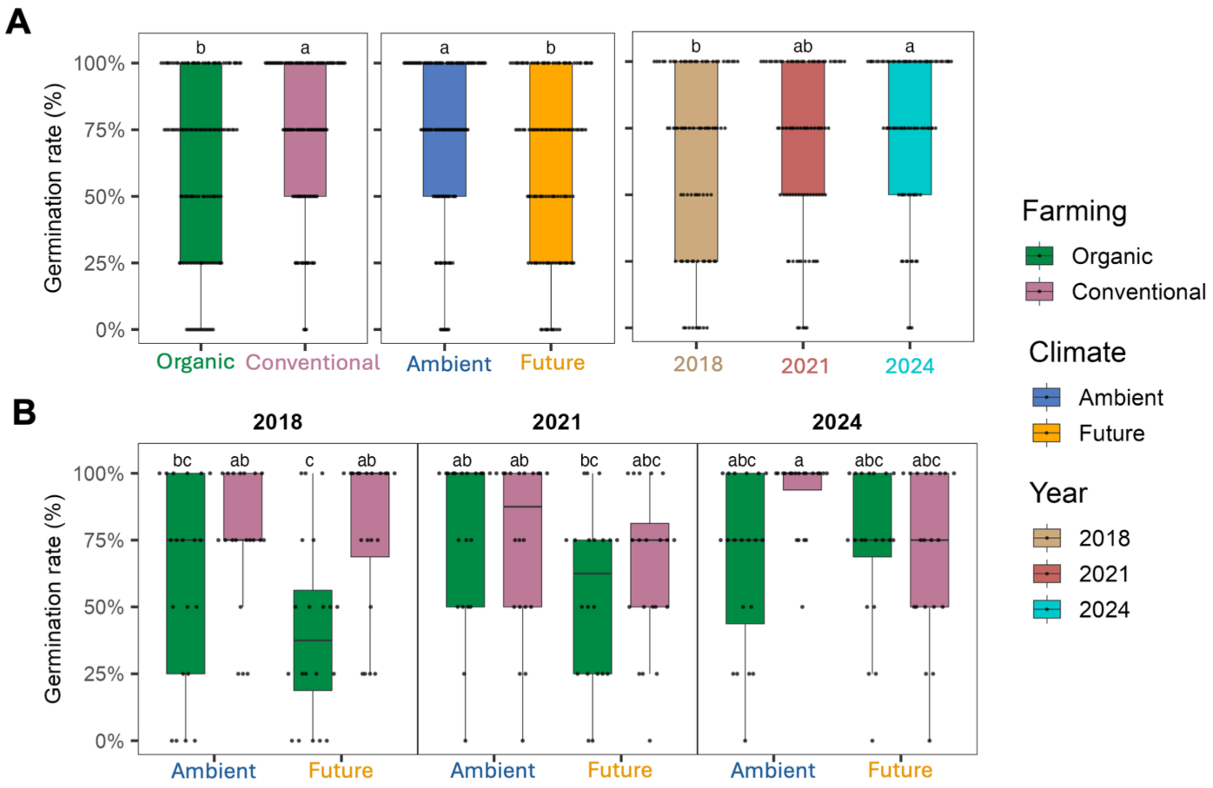


Figure S6: Germination rate on Day 6 by (A) farming, climate, harvest years and (B) significant interactions at controlled greenhouse experiment. Different letters in the panel denote statistically significant differences based on post hoc Tukey's test ($P$<0.05),

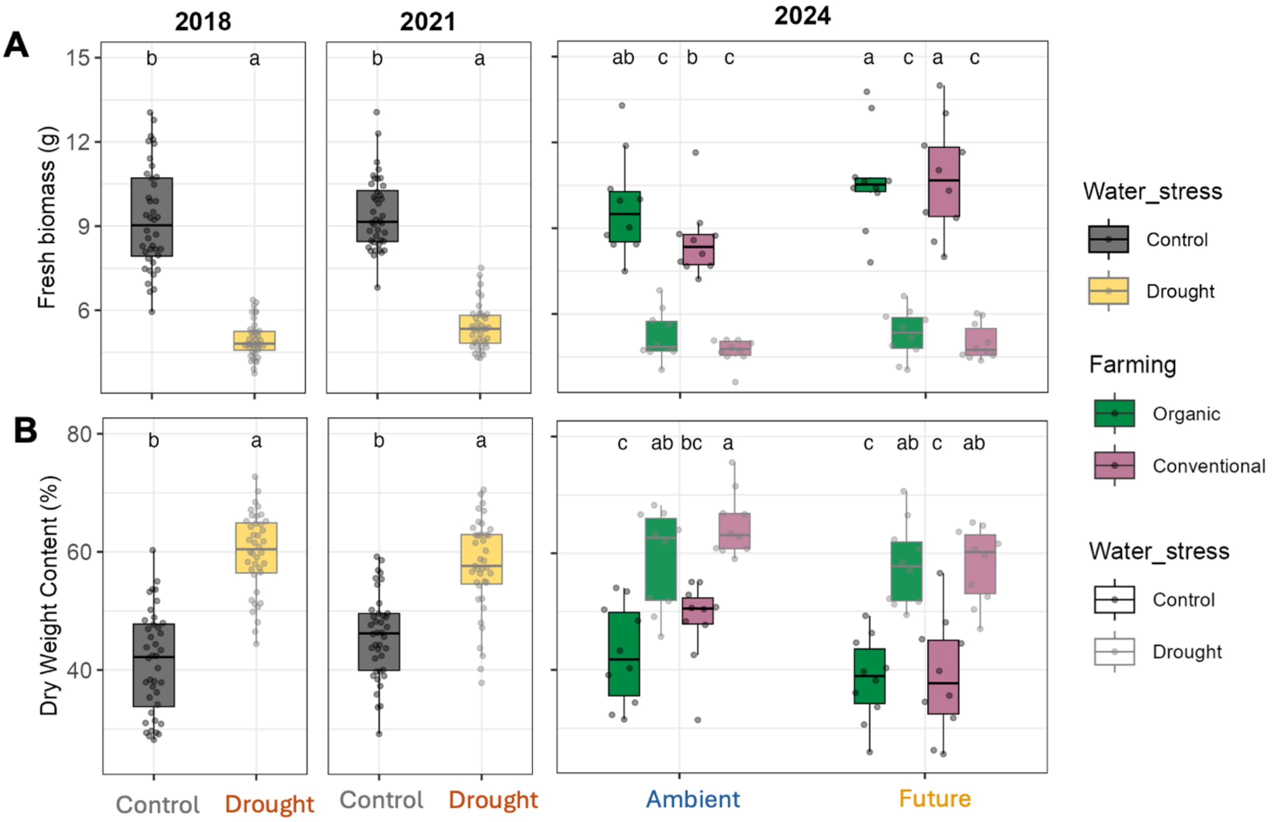


Figure S7: Physiological and microbial functional responses of offspring wheat plants under drought stress. (A) Fresh biomass and (B) dry weight content variations by water stress for seeds from the 2018 and 2021 harvest years and by farming and climate histories for seeds from 2024. Different letters in the panel denote statistically significant differences based on post hoc Tukey's test (*P*<0.05), whereas "ns" denotes non-significant interactions.

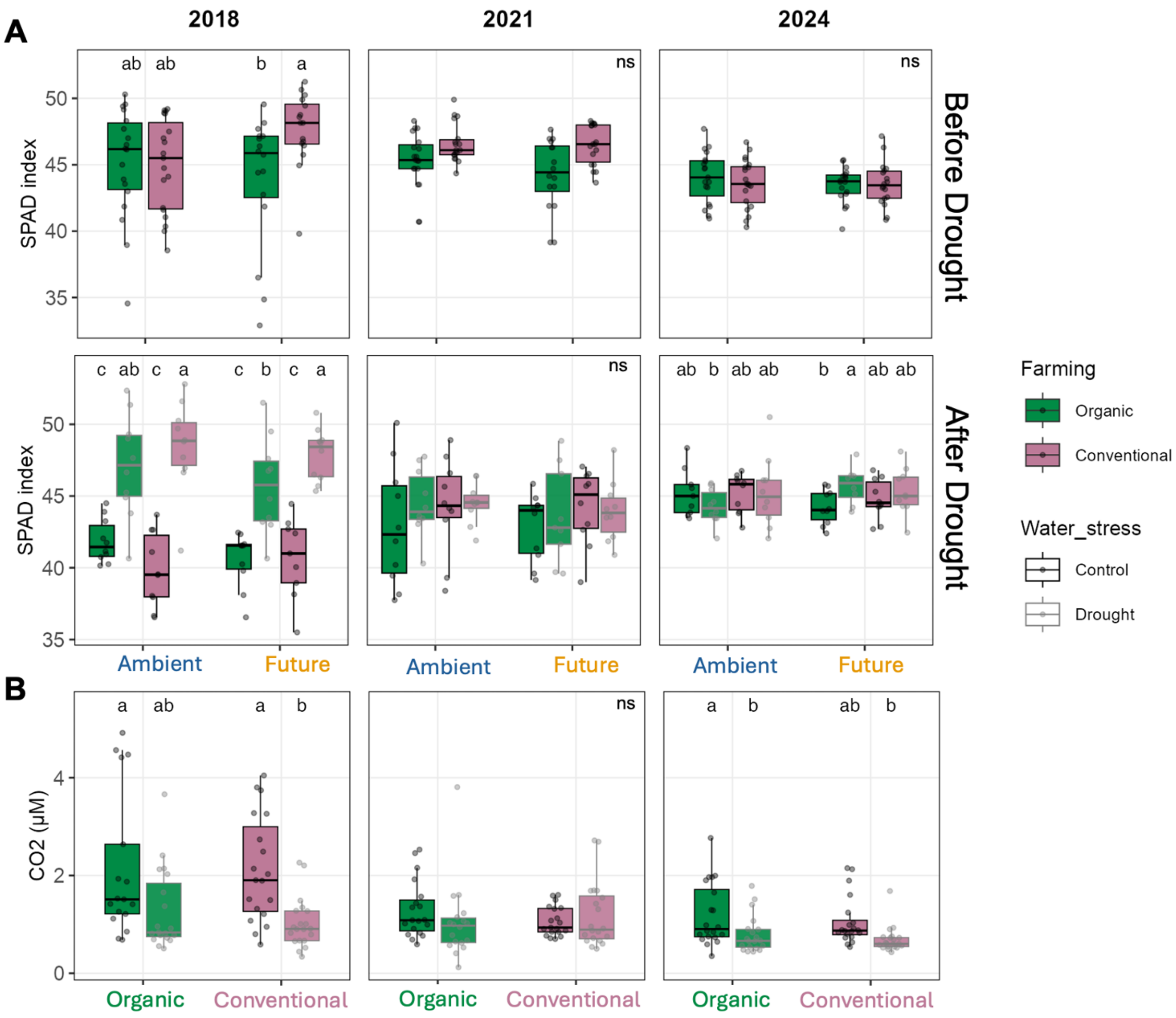


Figure S8: Physiological and microbial functional responses of offspring wheat plants under drought stress. (A) Variations in chlorophyll content (SPAD Index) before and after drought exposure on plants originating from seed with distinct farming and climatic history which were harvested in 2018, 2021, and 2024. (B) Microbial carbon respiration in the rhizosphere by farming history. Different letters in the panel denote statistically significant differences based on post hoc Tukey's test ($P<0.05$), whereas "ns" denotes non-significant interactions.

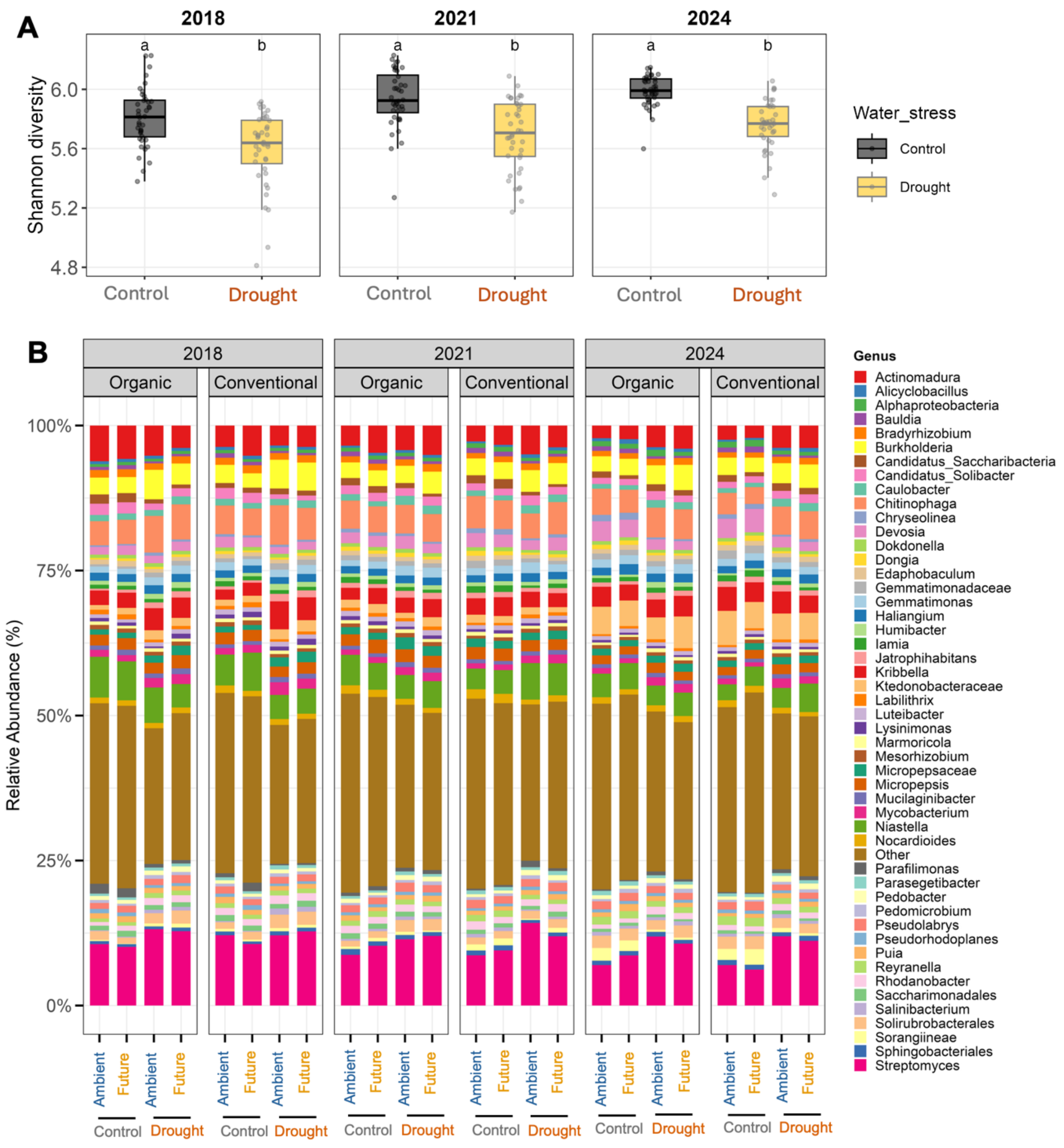


Figure S9: Rhizosphere bacterial diversity of the offspring wheat generation under contrasting seed microbial legacies. (A) Bacterial Shannon diversity of rhizosphere bacterial communities across harvest years (2018, 2021, and 2024) under organic and conventional farming legacies of seeds. Different letters denote statistically significant differences based on post hoc Tukey's test (*P*<0.05). (B) Relative abundance of top 50 most dominant bacterial genera detected in drought-affected rhizosphere across contrasting farming (organic vs. conventional) and climate (ambient vs. future) legacies of seeds harvested in 2018, 2021 and 2024.

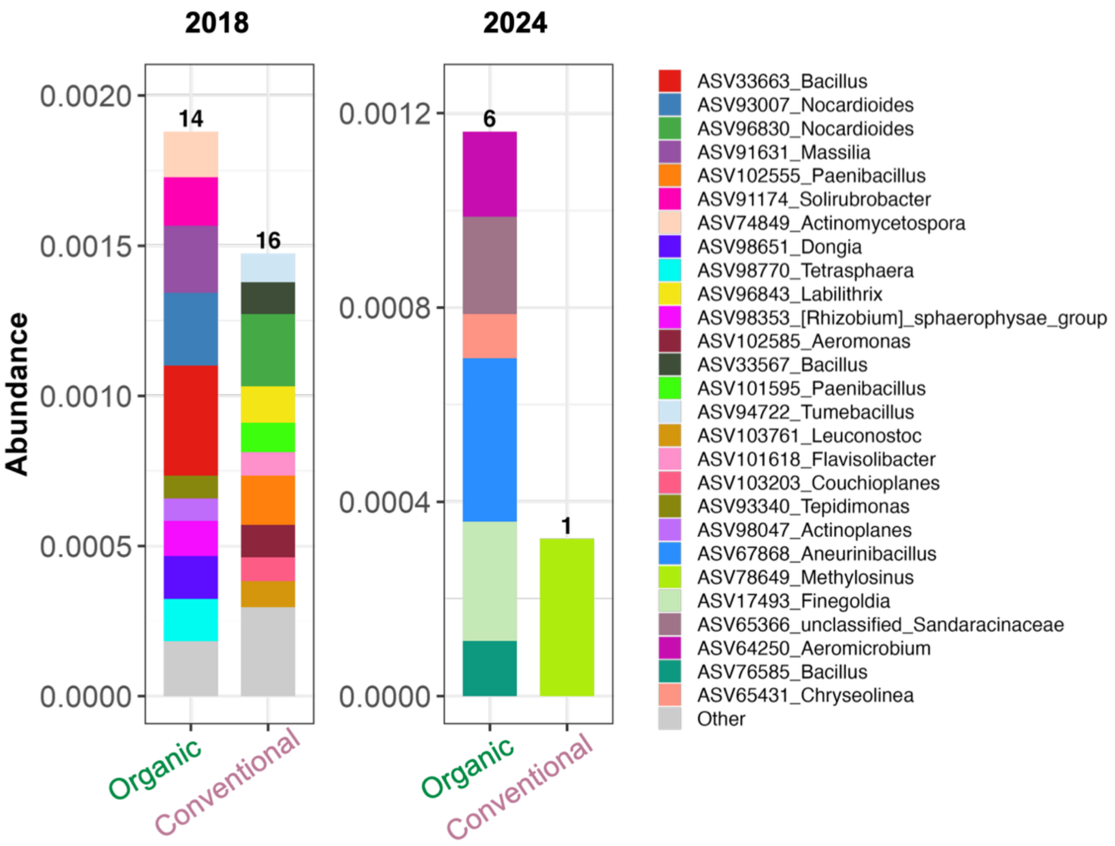


Figure S10. Abundance of year-specific unique bacterial ASVs detected in drought affected rhizosphere of offspring generation plants that originated from seed epiphytic and endophytic microbiomes under organic and conventional farming legacies across harvest years.

### Tables

**Table S1.** LMER results for the impact of Year, Month, and their interaction on daily air temperature and total precipitation. Data are presented for the full year and the growing season

(April–September). Significant p-values (< 0.05) are highlighted in bold. (F = f-value; P = P-value).

| **Factors** | **Temperature** | **Temperature (Growing season)** | **Precipitation** | **Precipitation (Growing season)** |
|---|---|---|---|---|
| *Month* | F = 57963.36 | F = 24780.74 | F = 20.308 | F = 17.826 |
| | **P < 0.001** | **P < 0.001** | **P < 0.001** | **P < 0.001** |
| *Year* | F = 562.66 | F = 633.33 | F = 12.008 | F = 11.143 |
| | **P < 0.001** | **P < 0.001** | **P < 0.001** | **P < 0.001** |
| *Month × Year* | F = 290.03 | F = 261.78 | F = 7.919 | F = 8.270 |
| | **P < 0.001** | **P < 0.001** | **P < 0.001** | **P < 0.001** |

*Bold values indicate statistical significance (P < 0.05).

**Table S2.** ANOVA test results for the effects of Farming, Climate, Year of harvest, and their interactions on dry weight of seeds from GCEF field (F = f-value; P = P-value).

| **Factors** | **Seed dry weight** |
|---|---|
| *Farming* | F = 54.64 |
| | **P < 0.001** |
| *Climate* | F = 7.021 |
| | **P = 0.003** |
| *Year* | F = 69.08 |
| | **P < 0.001** |
| *Farming × Climate* | F = 0.691 |
| | P = 0.411 |
| *Farming × Year* | F = 33.15 |
| | **P < 0.001** |
| *Climate × Year* | F = 4.829 |
| | **P = 0.013** |
| *Farming × Climate × Year* | F = 11.37 |
| | **P < 0.001** |

* Farming refers to the history of seed microbes from conventional and organic agriculture.
Climate refers to history of seed microbes from ambient and future climate under each farming.
Bold values indicate statistical significance (P < 0.05).

**Table S3.** ANOVA test results for the year wise-effects of Farming, Climate, and their interactions on dry weight of seeds from GCEF field (F = f-value; P = P-value).

| **Factors** | **2018** | **2021** | **2024** |
|---|---|---|---|
| *Farming* | F = 33.60 | F = 0.009 | F = 56.90 |
| | **P < 0.001** | P = 0.925 | **P < 0.001** |

| | | | |
|---|---|---|---|
| *Climate* | F = 31.29 | F = 0.028 | F = 1.500 |
| | **P = 0.001** | P = 0.870 | P = 0.256 |
| *Farming × Climate* | F = 32.74 | F = 1.621 | F = 9.152 |
| | **P < 0.001** | P = 0.221 | **P = 0.016** |

* Farming refers to the history of seed microbes from conventional and organic agriculture.
Climate refers to history of seed microbes from ambient and future climate under each farming.
Bold values indicate statistical significance (P < 0.05).

**Table S4.** ANOVA test results for the effects of Farming, Climate, harvest year, and their interactions on Shannon diversity of seed-associated bacteria (F = f-value; P = P-value).

| **Factors** | **Epiphyte** | **Endophyte** |
|---|---|---|
| *Farming* | F = 2.712 | F = 6.961 |
| | P = 0.106 | **P = 0.011** |
| *Climate* | F = 6.120 | F = 1.495 |
| | **P = 0.017** | P = 0.227 |
| *Year* | F = 32.225 | F = 8.855 |
| | **P < 0.001** | **P = 0.001** |
| *Farming × Climate* | F = 0.021 | F = 0.007 |
| | P = 0.886 | P = 0.934 |
| *Farming × Year* | F = 0.112 | F = 1.505 |
| | P = 0.894 | P = 0.232 |
| *Climate × Year* | F = 5.798 | F = 2.335 |
| | **P = 0.006** | P = 0.108 |
| *Farming × Climate × Year* | F = 1.471 | F = 0.435 |
| | P = 0.240 | P = 0.650 |

* Farming refers to the history of seed microbes from conventional and organic agriculture.
Climate refers to history of seed microbes from ambient and future climate under each farming.
Bold values indicate statistical significance (P < 0.05).

**Table S5.** ANOVA test results for the year wise-effects of Farming, Climate, and their interactions on Shannon diversity of seed-associated bacteria (F = f-value; P = P-value).

| **Factors** | **Epiphyte** | | | **Endophyte** | | |
|---|---|---|---|---|---|---|
| | **2018** | **2021** | **2024** | **2018** | **2021** | **2024** |
| *Farming* | F = 1.064 | F = 0.633 | F = 1.639 | F = 6.001 | F = 6.375 | F = 0.103 |
| | P = 0.318 | P = 0.438 | P = 0.219 | **P = 0.026** | **P = 0.023** | P = 0.752 |
| *Climate* | F = 0.735 | F = 2.893 | F = 43.12 | F = 0.086 | F = 0.000 | F = 4.379 |

| | | | | | | |
|---|---|---|---|---|---|---|
| | P = 0.404 | P = 0.108 | **P = 0.000** | P = 0.773 | P = 0.996 | P = 0.053 |
| *Farming × Climate* | F = 1.311 | F = 0.484 | F = 0.891 | F = 0.097 | F = 2.213 | F = 0.067 |
| | P = 0.269 | P = 0.496 | P = 0.359 | P = 0.760 | P = 0.156 | P = 0.799 |

* Farming refers to the history of seed microbes from conventional and organic agriculture.
Climate refers to history of seed microbes from ambient and future climate under each farming.
Bold values indicate statistical significance (P < 0.05).

**Table S6.** PERMANOVA test results for the effects of Farming, Climate, Year of harvest, and their interactions on community structure of seed-associated bacteria (F = f-value; P = P-value).

| Factors | Epiphyte | Endophyte |
|---|---|---|
| *Farming* | F = 0.734 | F = 1.685 |
| | P = 0.512 | P = 0.149 |
| *Climate* | F = 6.350 | F = 0.989 |
| | **P = 0.002** | P = 0.373 |
| *Year* | F = 45.30 | F = 10.35 |
| | **P = 0.001** | **P = 0.001** |
| *Farming × Climate* | F = 1.231 | F = 0.674 |
| | P = 0.268 | P = 0.586 |
| *Farming × Year* | F = 1.075 | F = 0.672 |
| | P = 0.300 | P = 0.727 |
| *Climate × Year* | F = 2.464 | F = 0.958 |
| | **P = 0.024** | P = 0.427 |
| *Farming × Climate × Year* | F = 1.085 | F = 0.472 |
| | P = 0.309 | P = 0.923 |

* Farming refers to the history of seed microbes from conventional and organic agriculture.
Climate refers to history of seed microbes from ambient and future climate under each farming.
Bold values indicate statistical significance (P < 0.05).

**Table S7.** PERMANOVA test results for the year wise-effects of Farming, Climate, and their interactions on community structure of seed-associated bacteria (F = f-value; P = P-value).

| Factors | Epiphyte | | | Endophyte | | |
|---|---|---|---|---|---|---|
| | 2018 | 2021 | 2024 | 2018 | 2021 | 2024 |
| *Farming* | F = 1.015 | F = 1.153 | F = 0.814 | F = 5.316 | F = 1.877 | F = 0.677 |
| | P = 0.422 | P = 0.297 | P = 0.597 | **P = 0.004** | P = 0.111 | P = 0.524 |
| *Climate* | F = 1.067 | F = 3.232 | F = 4.973 | F = 0.274 | F = 1.336 | F = 0.351 |
| | P = 0.325 | **P = 0.006** | **P = 0.001** | P = 0.957 | P = 0.218 | P = 0.719 |
| *Farming × Climate* | F = 2.012 | F = 0.776 | F = 1.138 | F = 0.528 | F = 0.538 | F = 0.207 |
| | **P = 0.021** | P = 0.656 | P = 0.285 | P = 0.710 | P = 0.820 | P = 0.918 |

* Farming refers to the history of seed microbes from conventional and organic agriculture.
Climate refers to history of seed microbes from ambient and future climate under each farming.

Bold values indicate statistical significance (P < 0.05).

**Table S10.** Generalized linear mixed model results for the effects of Farming, Climate, harvest years, and their interactions on germination at day 6 ($\chi^2$ = chi-square value; P = P-value).

| Factors | Germination |
|---|---|
| *Farming* | $\chi^2$ = 20.569 |
| | **P < 0.001** |
| *Climate* | $\chi^2$ = 12.912 |
| | **P < 0.001** |
| *Year* | $\chi^2$ = 11.913 |
| | **P = 0.003** |
| *Farming × Climate* | $\chi^2$ = 0.050 |
| | P = 0.822 |
| *Farming × Year* | $\chi^2$ = 9.743 |
| | **P = 0.008** |
| *Climate × Year* | $\chi^2$ = 1.460 |
| | P = 0.482 |
| *Farming × Climate × Year* | $\chi^2$ = 16.196 |
| | **P < 0.001** |

* Farming (df = 1) refers to the history of seed microbes from conventional and organic agriculture.
Climate (df = 1) refers to history of seed microbes from ambient and future climate under each farming.
Bold values indicate statistical significance (P < 0.05).

**Table S11.** ANOVA test results for the yearwise-effects of Farming, Climate, Water Stress, and their interactions on Plant height. (F = f-value; P = P-value).

| Factors | 2018 | | 2021 | | 2024 | |
|---|---|---|---|---|---|---|
| | Before | After | Before | After | Before | After |
| $R^2$ | **0.131** | **0.219** | **0.08** | **0.367** | **0.121** | **0.332** |
| *Farming* | F = 9.963 | F = 6.742 | F = 1.049 | F = 6.623 | F = 4.875 | F = 3.011 |
| | **P = 0.002** | **P = 0.011** | P = 0.309 | **P = 0.012** | **P = 0.030** | P = 0.087 |
| *Climate* | F = 1.146 | F = 0.192 | F = 4.053 | F = 2.071 | F = 3.996 | F = 4.471 |
| | P = 0.288 | P = 0.662 | **P = 0.048** | P = 0.154 | **P = 0.049** | **P = 0.038** |
| *Water Stress* | — | F = 9.497 | — | F = 26.942 | — | F = 26.552 |
| | — | **P = 0.003** | — | **P = $1.86 \times 10^{-6}$** | — | **P = $2.16 \times 10^{-6}$** |
| *Farming × Climate* | F = 0.346 | F = 0.127 | F = 1.484 | F = 3.586 | F = 1.630 | F = 0 |
| | P = 0.558 | P = 0.723 | P = 0.227 | P = 0.062 | P = 0.206 | P = 0.986 |

| | | | | | | |
|---|---|---|---|---|---|---|
| *Farming × Water Stress* | — | F = 1.925 | — | F = 0.007 | — | F = 0.04 |
| | — | P = 0.17 | — | P = 0.935 | — | P = 0.842 |
| *Climate × Water Stress* | — | F = 0.403 | — | F = 1.218 | — | F = 1.66 |
| | — | P = 0.527 | — | P = 0.273 | — | P = 0.202 |
| *Farming × Climate × Water Stress* | — | F = 1.309 | — | F = 1.279 | — | F = 0.09 |
| | — | P = 0.256 | — | P = 0.262 | — | P = 0.764 |

* Farming refers to the history of seed microbes from conventional and organic agriculture.
Climate refers to history of seed microbes from ambient and future climate under each farming.
Bold values indicate statistical significance (P < 0.05).

**Table S12.** ANOVA test results for the yearwise-effects of Farming, Climate, Water Stress, and their interactions on fresh biomass and dry weight content. (F = f-value; P = P-value).

| **Factors** | **Fresh Biomass** | | | **Dry Weight Content** | | |
|---|---|---|---|---|---|---|
| | **2018** | **2021** | **2024** | **2018** | **2021** | **2024** |
| *Farming* | F = 1.392 | F = 0.025 | F = 2.886 | F = 3.714 | F = 0.353 | F = 3.047 |
| | P = 0.242 | P = 0.875 | P = 0.094 | P = 0.058 | P = 0.554 | P = 0.085 |
| *Climate* | F = 3.965 | F = 1.128 | F = 8.921 | F = 2.561 | F = 1.693 | F = 10.111 |
| | P = 0.050 | P = 0.292 | **P = 0.004** | P = 0.114 | P = 0.197 | **P = 0.002** |
| *Water Stress* | F = 204.363 | F = 267.020 | F = 280.989 | F = 114.042 | F = 47.055 | F = 109.843 |
| | **P = 0.000** | **P = 0.000** | **P = 0.000** | **P = 0.000** | **P = 0.000** | **P = 0.000** |
| *Farming × Climate* | F = 0.702 | F = 0.267 | F = 1.695 | F = 0.214 | F = 0.530 | F = 2.721 |
| | P = 0.405 | P = 0.607 | P = 0.197 | P = 0.645 | P = 0.469 | P = 0.103 |
| *Farming × Water Stress* | F = 0.332 | F = 0.078 | F = 0.097 | F = 0.146 | F = 0.030 | F = 0.002 |
| | P = 0.566 | P = 0.781 | P = 0.756 | P = 0.704 | P = 0.864 | P = 0.965 |
| *Climate × Water Stress* | F = 2.595 | F = 0.486 | F = 5.544 | F = 0.588 | F = 0.166 | F = 0.820 |
| | P = 0.112 | P = 0.488 | **P = 0.021** | P = 0.446 | P = 0.685 | P = 0.368 |
| *Farming × Climate × Water Stress* | F = 1.191 | F = 1.240 | F = 0.801 | F = 0.025 | F = 1.702 | F = 0.003 |
| | P = 0.279 | P = 0.269 | P = 0.374 | P = 0.875 | P = 0.196 | P = 0.957 |

* Farming refers to the history of seed microbes from conventional and organic agriculture.
Climate refers to history of seed microbes from ambient and future climate under each farming.
Bold values indicate statistical significance (P < 0.05).

**Table S13.** ANOVA test results for the yearwise-effects of Farming, Climate, Water Stress, and their interactions on SPAD index (F = f-value; P = P-value).

| **Factors** | **2018** | | **2021** | | **2024** | |
|---|---|---|---|---|---|---|
| | **Before** | **After** | **Before** | **After** | **Before** | **After** |
| *Farming* | F = 2.966 | F = 0.647 | F = 3.001 | F = 1.416 | F = 0.285 | F = 0.009 |
| | P = 0.090 | P = 0.424 | P = 0.088 | P = 0.238 | P = 0.595 | P = 0.927 |
| *Climate* | F = 0.688 | F = 0.690 | F = 0.196 | F = 0.390 | F = 0.536 | F = 0.413 |
| | P = 0.410 | P = 0.409 | P = 0.659 | P = 0.534 | P = 0.467 | P = 0.523 |
| *Water Stress* | — | F = 116.516 | — | F = 0.832 | — | F = 1.600 |
| | — | **P < 0.001** | — | P = 0.365 | — | P = 0.210 |
| *Farming × Climate* | F = 4.259 | F = 1.493 | F = 1.402 | F = 0.023 | F = 0.433 | F = 1.203 |
| | **P = 0.043** | P = 0.226 | P = 0.240 | P = 0.881 | P = 0.513 | P = 0.277 |
| *Farming × Water Stress* | — | F = 5.062 | — | F = 0.926 | — | F = 0.405 |
| | — | **P = 0.028** | — | P = 0.339 | — | P = 0.527 |
| *Climate × Water Stress* | — | F = 0.373 | — | F = 0.272 | — | F = 5.126 |

| | | | | | | |
|---|---|---|---|---|---|---|
| | — | P = 0.544 | — | P = 0.603 | — | **P = 0.027** |
| *Farming × Climate × Water Stress* | — | F = 0.323 | — | F = 0.011 | — | F = 3.059 |
| | — | P = 0.571 | — | P = 0.915 | — | P = 0.085 |

* Farming refers to the history of seed microbes from conventional and organic agriculture.
Climate refers to history of seed microbes from ambient and future climate under each farming.
Bold values indicate statistical significance (P < 0.05).

**Table S14.** ANOVA test results for the yearwise-effects of Farming, Climate, Water Stress, and their interactions on carbon respiration of rhizosphere microbes. (F = f-value; P = P-value).

| **Factors** | **2018** | **2021** | **2024** |
|---|---|---|---|
| *Farming* | F = 0.794 | F = 0.126 | F = 1.270 |
| | P = 0.376 | P = 0.723 | P = 0.263 |
| *Climate* | F = 2.689 | F = 0.001 | F = 1.692 |
| | P = 0.106 | P = 0.972 | P = 0.198 |
| *Water Stress* | F = 10.83 | F = 0.193 | F = 13.27 |
| | **P = 0.002** | P = 0.662 | **P = 0.001** |
| *Farming × Climate* | F = 0.687 | F = 0.589 | F = 0.244 |
| | P = 0.411 | P = 0.445 | P = 0.623 |
| *Farming × Water Stress* | F = 5.933 | F = 1.721 | F = 0.020 |
| | **P = 0.018** | P = 0.194 | P = 0.887 |
| *Climate × Water Stress* | F = 3.041 | F = 0.546 | F = 0.060 |
| | P = 0.086 | P = 0.462 | P = 0.807 |
| *Farming × Climate × Water Stress* | F = 0.262 | F = 0.008 | F = 0.026 |
| | P = 0.610 | P = 0.928 | P = 0.872 |

* Farming refers to the history of seed microbes from conventional and organic agriculture.
Climate refers to history of seed microbes from ambient and future climate under each farming.
Bold values indicate statistical significance (P < 0.05).
